# Bidirectional Extracellular Vesicle-Mediated Maternal-Embryonic Exchange in the Lecithotrophic Teleost Guppy

**DOI:** 10.64898/2026.04.25.717372

**Authors:** Junki Yoshida, Kazuko Uchida, Makoto Kuwahara, Eiichi Hondo, Natsuko Kawano, Atsuo Iida

**Affiliations:** Department of Life Sciences, School of Agriculture, Meiji University, Tama-ku, Kawasaki, Kanagawa, Japan; Organization for the Strategic Coordination of Research and Intellectual Properties, Meiji University, Tama-ku, Kawasaki, Kanagawa, Japan; Institute of Materials and Systems for Sustainability, Nagoya University, Furo-cho, Chikusa-ku, Nagoya, 464–8601, Japan; Department of Animal Sciences, Graduate School of Bioagricultural Sciences, Nagoya University, Furo-cho, Chikusa-ku, Nagoya, 464–8601, Japan

**Author notes:** Paste corresponding author name(s) here. PNAS requires the corresponding author to provide an ORCID identifier at submission and strongly encourages all authors to use an ORCID ID. Do not include ORCIDs in the manuscript file; individual authors must link their ORCID account to their PNAS profile at www.pnascentral.org. For proper authentication, authors are not permitted to add ORCIDs on proofs. Learn more or register for ORCID. Author Contributions: J.Y. designed the study; J.Y. and K.U. performed the experiments; M.K., E.H., and N.K. provided new reagents and analytic tools; J.Y. analyzed the data; and J.Y. and A.I, wrote the original draft; K.U., M.K., E.H., and N.K., reviewed and edited the original draft. All authors have read and agreed to the published version of the manuscript. Competing Interest Statement: The authors declare no conflicts of interest. Classification: BIOLOGICAL SCIENCES, Developmental Biology; Physiology.

**Keywords:** Maternal-fetal interaction, Poeciliid fish, Extracellular vesicles, Viviparity, Maternal investment

## Abstract

The placenta is defined as an organ that mediates the exchanges of nutrients, hormones, and other substances between the mother and embryo in viviparous animals. Its structure is diverse due to interspecific differences in fetal tissues and variation in the forms of maternal-fetal interfaces. Matrotrophic poeciliid teleosts, in which the embryo develops within the maternal ovarian follicle, possess functional placentas formed from maternal follicular and embryonic tissues. However, the physiological mechanisms underlying substrate exchange between mother and embryo remain unclear. Furthermore, although similar embryonic-maternal interfaces are observed in lecithotrophic poeciliids, it is unknown whether nutrient exchange occurs in these species. Therefore, this study investigated whether substance exchange occurs between the mother and embryo in the lecithotrophic teleost guppy (*Poecilia reticulata*) and identified the underlying physiological mechanisms. Histological analysis revealed that guppies have embryonic-maternal interfaces consisting of the maternal follicle and the embryonic yolk sac and pericardial sac. Additionally, tracking of 2,000 kDa fluorescein isothiocyanate-dextran injected into pregnant guppies confirmed its transport from mother to embryo. Immunofluorescence staining and electron microscopy revealed that substance transport from mother to embryo occurs via extracellular vesicles. Moreover, immunofluorescence staining and pharmacological experiments revealed exosome transport from embryo to mother. This study demonstrates that lecithotrophic guppies possess a functional placenta that mediates maternal-embryonic substrate transfer via extracellular vesicles. These findings provide fundamental insight into the evolution of placental strategies within the *Poeciliidae* family.

**Significance Statement:** Viviparity, in which embryos develop within the maternal body, has evolved independently across diverse animal lineages. In lecithotrophic viviparity, embryos are thought to rely primarily on yolk-derived nutrients, with maternal–fetal exchange limited to small molecules such as gases. Here, using macromolecular tracer experiments and ultrastructural analyses in guppies, we show that large macromolecules (2000 kDa) are exchanged bidirectionally between mother and fetus via extracellular vesicles, despite the absence of direct tissue attachment. These findings challenge the conventional view of lecithotrophic viviparity and reveal a previously unrecognized mechanism of maternal–fetal communication. Our results suggest that extracellular vesicle–mediated exchange may represent a widespread and evolutionarily conserved strategy for maternal–fetal interaction across viviparous animals.

## Introduction

Sexual reproduction in animals is broadly classified into oviparity, in which eggs are laid externally, and viviparity, in which embryos develop and are born within the maternal body. Viviparity can be further divided into lecithotrophic viviparity, in which embryonic development relies primarily on yolk-derived nutrients, and matrotrophic viviparity, in which the mother supplies nutrients to the developing embryos during gestation. Matrotrophic viviparity encompasses a wide range of nutritional strategies. In placentotrophy, maternal and fetal tissues form specialized interfaces, such as the placenta, to facilitate nutrient transfer. In contrast, histotrophy, which has been observed in various taxa, including certain stingrays and tsetse flies, involves the secretion of nutrients from the maternal reproductive tract that are ingested by the embryo. Additional reproductive strategies include embryophagy, in which developing embryos consume siblings, and oophagy, in which unfertilized eggs serve as a nutritional source(1). Together, these reproductive modes illustrate the remarkable diversity of maternal nutritional investment during embryonic development.

The placenta is broadly regarded as a maternal–embryonic interface that facilitates physiological exchange between mother and offspring during gestation, although its precise definition varies across taxa and disciplines. Owing to the diversity of tissues contributing to placental structures, this organ cannot be defined solely by developmental homology. For example, in humans and mice, the placenta is formed from the fetal mesoderm-derived chorion in association with the maternal endometrium(2, 3). In contrast, during early eutherian development and throughout gestation in marsupials, an endoderm-derived choriovitelline placenta serves as the primary placental structure(4, 5). In cartilaginous fishes, including certain sharks, placentation involves the fetal yolk sac and the maternal uterine epithelium. Moreover, in many eutherians, marsupials, and *Mabuya* lizards, both the choriovitelline membrane and the yolk sac contribute to placental function(6–8). Even among mammals, placental morphology varies substantially: in humans and mice, fetal trophoblasts invade the maternal decidua to form a hemochorial placenta, whereas in cattle and horses, trophoblasts adhere to but do not invade maternal tissues, thereby forming an epitheliochorial placenta(3). Despite this structural diversity, placentas can be broadly regarded as functional maternal–fetal interfaces that enable material exchange during pregnancy.

Fish placenta for nutrient transfer systems have also evolved in teleost fishes. In goodeid fishes, post-hatching embryos absorb maternally secreted nutrients, such as glucose and vitellogenin, via hindgut-derived structures known as trophotaeniae through endocytosis(9–12). In the family *Jenynsia*, nutrient transfer is thought to occur via projections from the maternal ovary and the gills of developing embryos(13). These structures are often referred to as pseudoplacentas in the literature, highlighting their functional similarity to placentas despite their distinct developmental origins.

The family *Poeciliidae*, belonging to the order *Cyprinodontiformes*, comprises predominantly viviparous species in which fertilization and embryonic development occur within the ovarian follicles. In this family, the matrotrophic index (MI; dry mass of offspring at birth relative to the dry mass of the egg) varies widely among species, and comparative analyses have revealed multiple independent transitions between lecithotrophy and matrotrophy(14). Embryos of poeciliid fishes possess extraembryonic membrane tissues, including the yolk sac and a pericardial sac that extends over the head and heart. These membranes are richly vascularized and lie in close proximity to the maternal ovarian follicle, thereby forming the maternal–embryonic interface. In matrotrophic poeciliids, the ovarian follicles undergo pronounced morphological changes during pregnancy, including thickening and the formation of villus(15). However, unlike mammalian placentas, maternal and embryonic tissues in poeciliids do not exhibit direct tissue adhesion, leaving the mechanisms of substance exchange between mother and embryos unresolved.

Guppies (*Poecilia reticulata*) are classified as lecithotrophic viviparous fish because the dry mass of eggs exceeds that of live-born offspring(16). Consequently, maternal nutritional investment has been assumed to occur prior to fertilization, with the maternal–embryonic interface functioning primarily in gas exchange and waste removal. However, in vitro culture studies have shown that fetal bovine serum is required for successful guppy embryo development and that embryonic survival and growth depend on its concentration(17), suggesting that guppy embryos may require exogenous substances beyond yolk reserves. Moreover, embryos of lecithotrophic poeciliids, including guppies, possess a pericardial sac that has been implicated in nutrient absorption in matrotrophic relatives(15, 18, 19). These observations raise the possibility that maternal–embryonic material exchange also occurs in lecithotrophic poeciliids.

In this study, we investigated whether substance exchange occurs between mother and embyos in guppies and examined the physiological mechanisms underlying this exchange. Our data reveal that bidirectional maternal–embryonic transport occurs via extracellular vesicles, providing new insight into the diversity of placental strategies and the evolution of maternal provisioning within the Poeciliidae.

## Results

### Structure of maternal-to-embryonic interface in pregnant guppies

Poeciliid embryos possess two extraembryonic membrane tissues containing blood vessels: the yolk sac and the pericardial sac (Fig. 1A and B). In guppies, the pericardial sac extends over the head region, and vascular networks are observed on both the pericardial sac and the yolk sac (Fig. 1A and B; Movies S1 and S2). The majority of these vessels form dense capillary plexuses, in addition to identifiable larger vessels, including portal veins surrounding the heart and paired vitelline arteries branching from the trunk (Fig. 1A).

**Figure 1.**
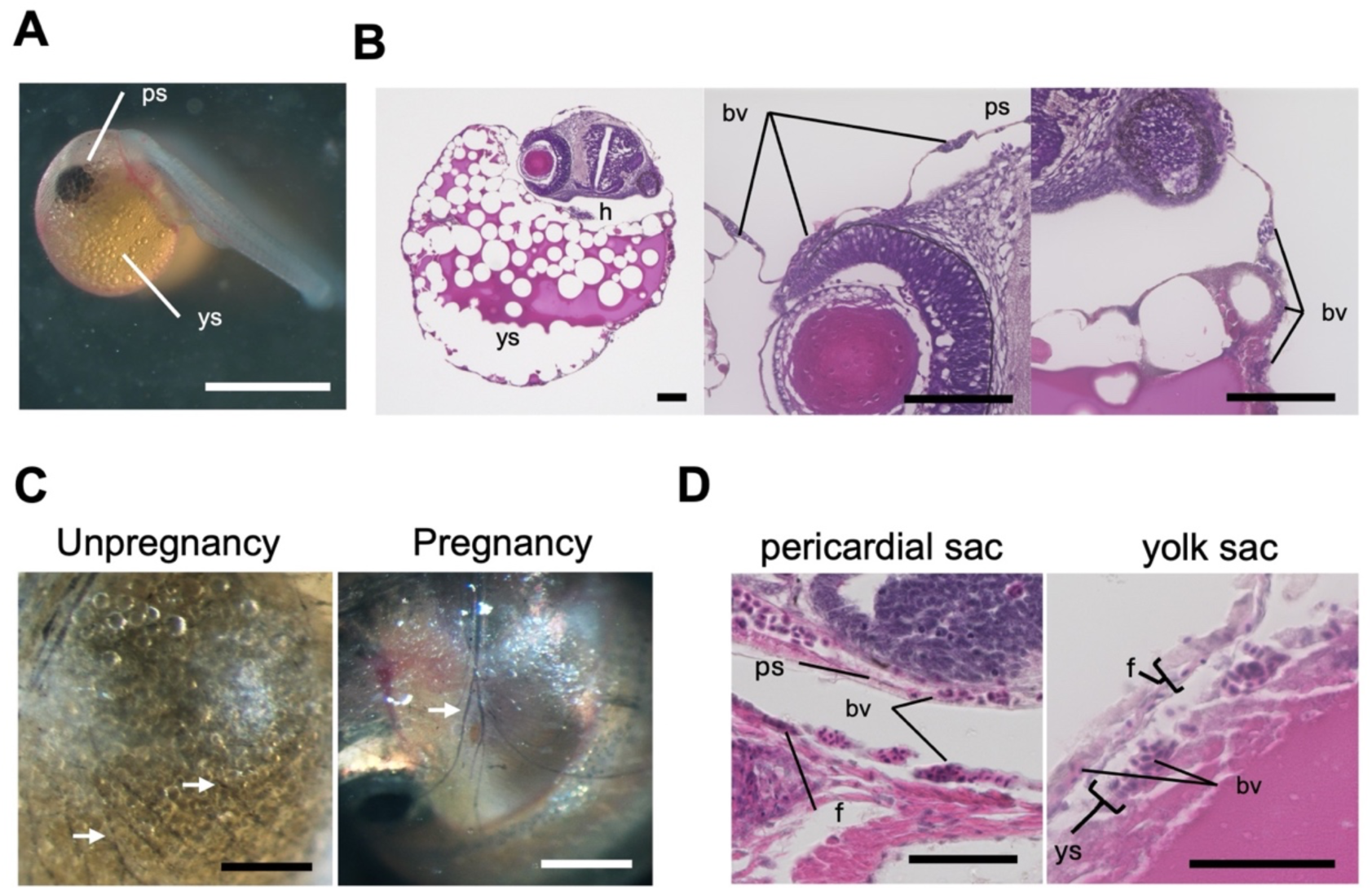
Placenta-like structure in guppies. (A) Photograph of a guppy embryo at Mousavi’s developmental stage pharyngula 29 (Ph29)(19). The white scale bar indicates 1 mm. ps: pericardial sac; ys: yolk sac. **(B)** Histological analysis of a stage Ph29 embryo. The scale bar indicates 100 µm. bv: blood vessel; h: heart; ps: pericardial sac; ys: yolk sac. **(C)** Follicular vasculature visualized by black ink injection in the nonpregnant ovary (left) and the late-pregnant ovary (right). The scale bar indicates 500 µm. **(D)** Histological analysis of the maternal–embryonic interface in pregnant guppies. The left panel shows the interface between the maternal follicle and the embryonic pericardial sac. The right panel shows the interface between the maternal follicle and the embryonic yolk sac. The scale bar indicates 50 µm. f: follicle; ps: pericardial sac; ys: yolk sac.

To examine maternal vasculature, black ink was injected into the abdominal cavity of females. Ink perfusion enabled visualization of capillary plexuses in the caudal fin within 1 h (*SI Appendix*, Fig. S1), indicating effective systemic perfusion. In nonpregnant ovaries (n = 6), follicular capillaries formed a dense mesh-like network surrounding each oocyte (Fig. 1C). In contrast, mid-gestation follicles (n = 7) exhibited a more organized vascular arrangement characterized by larger-caliber vessels giving rise to smaller branching capillaries (Fig. 1C).

To characterize the embryonic–maternal interface, anatomical and histological analyses were performed on mid-gestation follicles (Fig. 1D). Hematoxylin–eosin staining showed that the maternal follicular epithelium was closely apposed to the embryonic pericardial sac, yolk sac, and embryonic body. No continuous tissue fusion or interdigitation structures were observed at the light microscopic level. In addition, embryonic movements caused positional changes of the embryo within the maternal follicle (Movie S3).

### Maternal-to-embryonic transfer of FITC-dextran in pregnant guppies

To examine whether substances are transferred from mother to embryo during pregnancy, 2,000 kDa FITC-dextran or PBS was injected into the abdominal cavity of mid-gestation females, and fluorescence in their embryos was observed (Fig. 2A).

**Figure 2.**
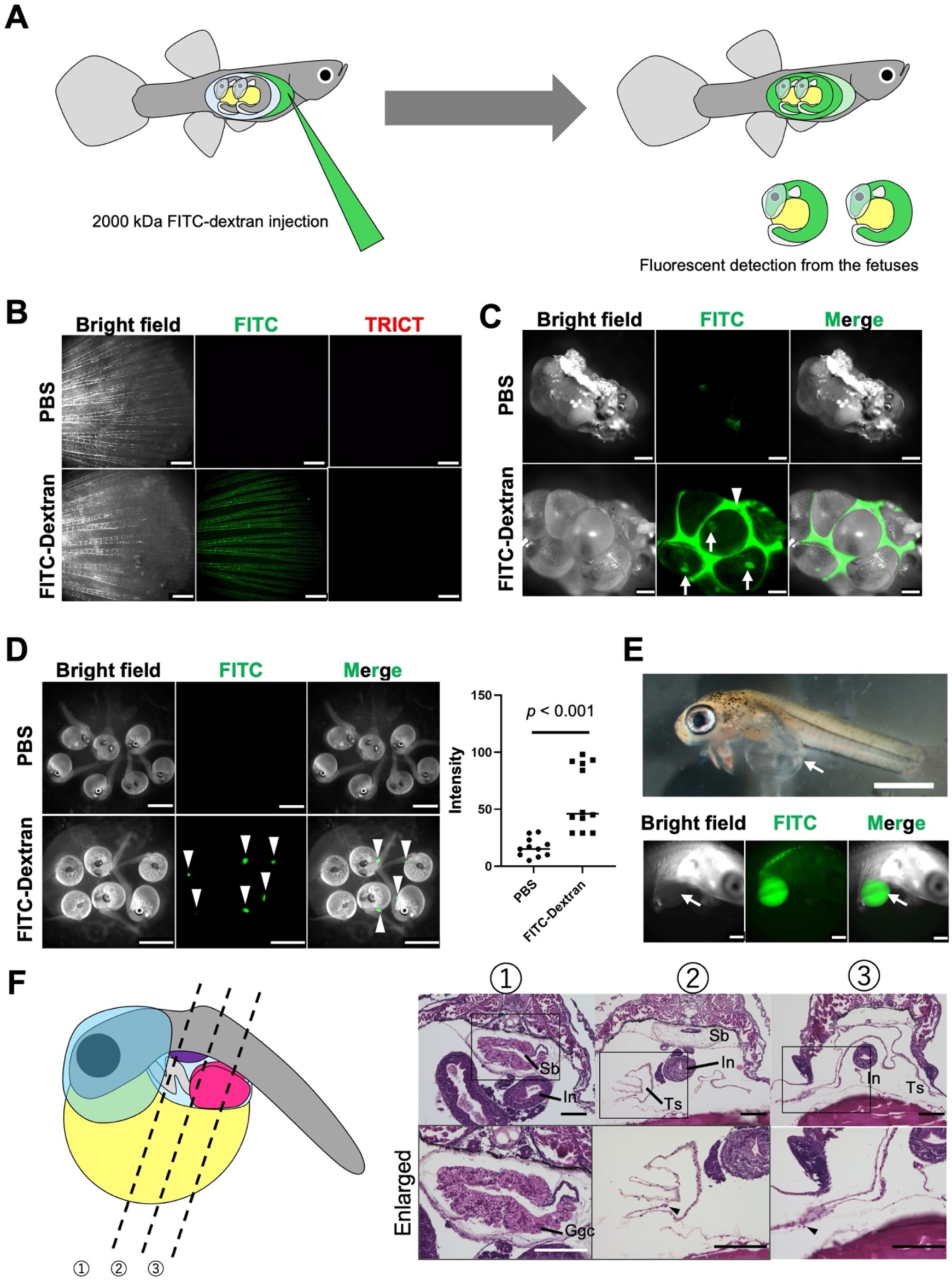
Maternal-to-embryonic transport of substances in pregnant guppies. **(A–F)** Experiments were performed to examine whether 2,000 kDa FITC–dextran injected into pregnant females was transported to the embryo. **(A)** Schematic illustration of the experimental design. Mid-gestation females were intraperitoneally injected with 2,000 kDa FITC-dextran or PBS, and FITC signals in the ovaries and embryos were examined 24 h after injection. **(B)** Fluorescence stereomicroscopic observation of the caudal fin of pregnant females 1 h after injection of PBS or FITC-dextran. The top row shows the caudal fin of a PBS-injected female, and the bottom row shows that of a FITC-dextran–injected female. Scale bar, 1 mm. **(C)** Fluorescence stereomicroscopic observation of ovaries from pregnant females 24 h after injection of PBS or FITC-dextran. The upper panel shows ovaries from a PBS-injected female, and the lower panel shows ovaries from a FITC-dextran–injected female. White arrows indicate FITC signals detected within follicles containing embryos, and white arrowheads indicate signals detected in the interfollicular space. Scale bar, 1 mm. **(D)** The left panels show fluorescence stereomicroscopic observation of embryos from pregnant females 24 h after injection of PBS or FITC-dextran. The top row shows embryos from PBS-injected females, and the bottom row shows embryos from FITC-dextran–injected females. White arrowheads indicate FITC signals detected within the embryo. Scale bar, 2 mm. The right panel shows quantification of the maximum fluorescence intensity detected in the fetuses shown in the left panels. An unpaired two-tailed Student’s *t*-test was conducted. **(E)** Accumulation of FITC–dextran in the trophic sac. The upper panel shows a stage Ph36 embryo observed under a stereomicroscope after removal of the yolk sac and yolk to expose the trophic sac. The lower panel shows a embryo from a FITC-dextran–injected female with the yolk removed to expose the trophic sac. White arrows indicate the trophic sac. Scale bars, 1 mm (upper) and 500 µm (lower). **(F)** Histological analysis of the trophic sac in a stage Ph29 embryo. The left schematic indicates the anatomical positions corresponding to the HE-stained sections shown on the right. The upper right panel shows the embryonic body cavity region, and the lower right panel shows a higher-magnification view of the boxed area in the upper panel. Sb: swim bladder; in: intestine; Ggc: gas gland cells; Ts: trophic sac. Scale bar, 50 µm.

Observation of caudal fin capillaries at 1 h and 24 h post-injection reveals fluorescent signals in FITC-dextran–injected females, whereas no fluorescence is detected in PBS-injected controls (Fig. 2B, *SI Appendix*, Fig. S2). The fluorescent signal is confined to vascular structures, and no extravascular diffusion is observed in the fin tissue at either time point.

Twenty-four hours after injection, fluorescent signals are detected in the interfollicular spaces and within ovarian follicles of FITC-dextran–injected females (Fig. 2C). In contrast, ovaries from PBS-injected females show no comparable fluorescence, aside from weak autofluorescence in septal tissues.

To assess whether embryonic tissues contain dextran, embryos were carefully separated from follicles and examined under fluorescence microscopy. No fluorescence is observed in embryos derived from PBS-injected females (Fig. 2D). In contrast, embryos from FITC-dextran–injected females exhibit distinct fluorescence within the body cavity (Fig. 2D). In addition, quantitative analysis of fluorescence intensity revealed that embryos from mothers injected with FITC-dextran showed significantly higher fluorescence intensity than embryos from PBS-injected negative controls.

Anatomical analysis reveals paired sac-like structures within the embryonic body cavity, hereafter referred to as trophic sacs (Fig. 2E). Fluorescence signals are localized within these trophic sacs.

Histological analysis of embryos at Mousavi’s developmental stage pharyngula 29 (Ph 29)(19) shows that the trophic sacs are anatomically distinct from the swim bladder (Fig. 2F). The swim bladder is located dorsally and contains gas gland cells, whereas the trophic sacs are positioned adjacent to the intestinal tract and lack gas gland cells. The trophic sacs extend bilaterally on the cephalic side of the intestine and fuse dorsally on the anal side.

To examine the role of the maternal follicle in embryonic uptake of FITC-dextran, embryos with the follicle intact and embryos after follicle removal were cultured in medium containing 1 mg/ml 2000 kDa FITC-dextran (*SI Appendix*, Fig. S3). In embryos with the follicle intact (n = 13), fluorescence was detected in the yolk and trophic sac but not in the mouth, gills, or digestive tract. In contrast, embryos cultured after follicle removal (n = 8) showed fluorescence in the mouth, gills, and digestive tract but not in the trophic sac. These results indicate that the route of FITC-dextran uptake differs depending on the presence of the maternal follicle.

### Maternal-to-embryonic transfer of clustered extracellular vesicles in pregnant guppies

The FITC-dextran injection experiments demonstrated that macromolecules are transported from the mother to the embryos. We therefore hypothesized that this maternal–embryonic transport may occur via extracellular vesicles.

To test this possibility, ultrathin sections of the maternal follicle and embryonic yolk sac were examined by electron microscopy to determine whether extracellular vesicle–like structures are present at the maternal–embryonic interface (Fig. 3A). Extracellular-Vesicle-like (EV-like) structures are observed within the follicular cavity adjacent to the embryo. These structures exhibit high electron density and range from approximately 100–300 nm in diameter. Clusters of such vesicle-like structures are observed in the follicular cavity.

**Figure 3.**
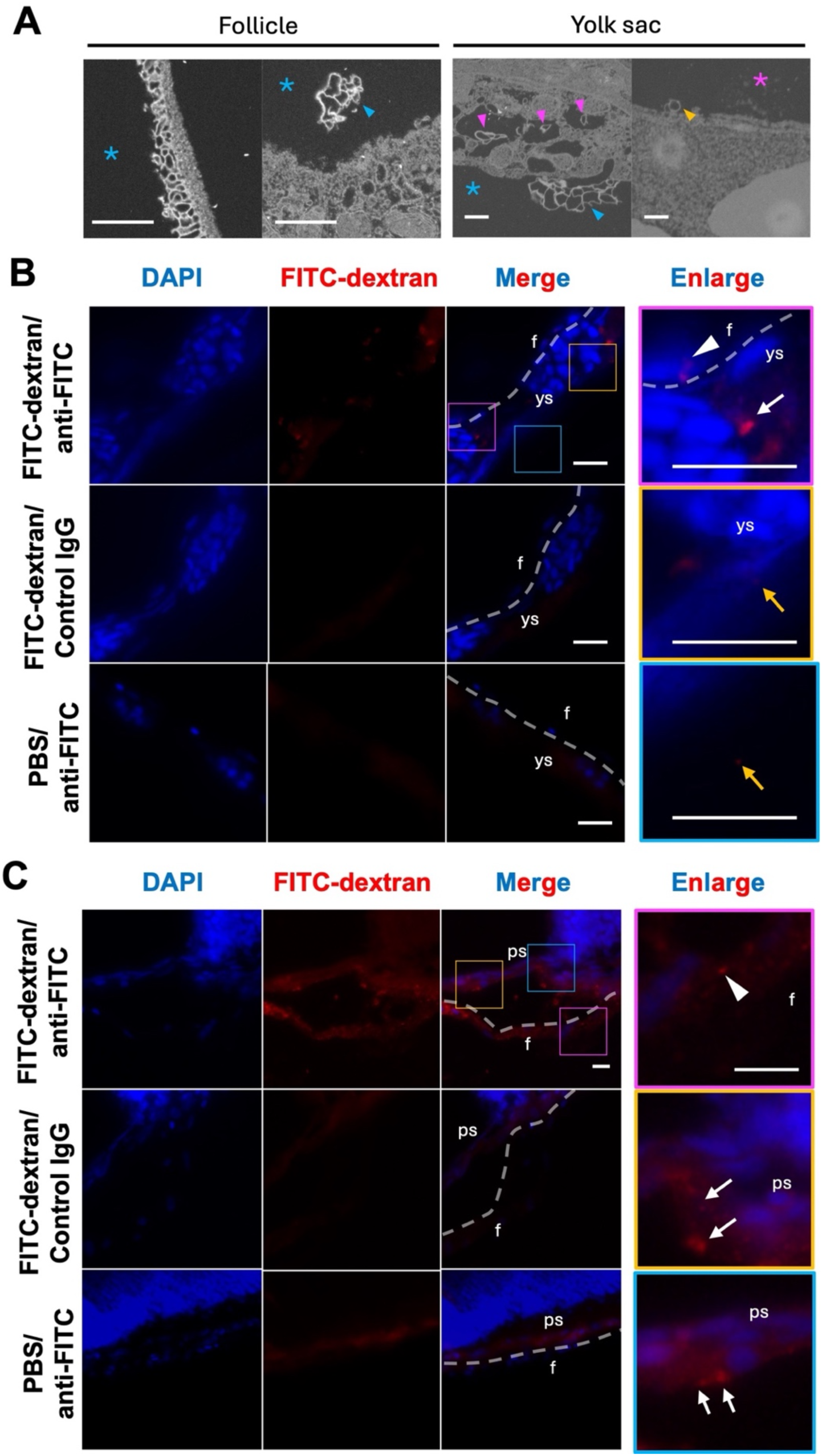
Maternal-to-embryonic substance transport via extracellular vesicles. **(A)** Scanning electron microscopy of ultrathin sections from a mid-gestation follicle and the enclosed embryonic yolk sac. The two left panels show ultrastructural features of the maternal follicle, and the two right panels show those of the embryonic yolk sac. Blue asterisks indicate the follicular cavity in which the follicle contacts the yolk sac, and magenta asterisks indicate the interior of the yolk sac where yolk accumulates. Blue arrowheads indicate clusters of extracellular vesicles secreted from the follicle. Magenta arrowheads indicate clusters of extracellular vesicles internalized by yolk sac cells via endocytosis. Yellow arrowheads indicate extracellular vesicles secreted from yolk sac cells into the yolk sac lumen. Scale bar, 500 nm. **(B, C)** Immunofluorescence staining was performed to examine whether 2,000 kDa FITC–dextran injected into pregnant females is directly transported from the maternal follicle to the embryonic yolk sac **(B)** or pericardial sac **(C)**. In each panel, the upper row (excluding the “Enlarged” column) shows immunofluorescence staining of embryos within follicles dissected from females injected with FITC-dextran, using an anti-FITC antibody as the primary antibody. The middle row shows staining of the same specimens using rabbit IgG as a primary antibody control. The bottom row shows immunofluorescence staining using an anti-FITC antibody on embryos within follicles dissected from females injected with PBS. “Enlarged” panels show magnified views of the merged FITC–dextran/anti-FITC signals, highlighting regions in which signal localization is confirmed. In **(B)**, the magenta panel indicates the follicle–yolk sac contact region, the yellow panel indicates the basal region of the yolk sac, and the blue panel indicates the interior of the yolk sac. In **(C)**, the magenta panel indicates the follicle, whereas the yellow and blue panels indicate the pericardial sac. Blue signals indicate DAPI-labeled nuclei, and red signals indicate FITC-dextran detected by immunofluorescence. White dotted lines indicate the boundary between the maternal follicle and the embryonic yolk sac or pericardial sac. White arrows indicate FITC-dextran signals detected in the maternal follicle, and white arrowheads indicate FITC-dextran signals detected in the yolk sac. Yellow arrows indicate FITC-dextran signals detected in the yolk sac lumen. f: maternal follicle; ys: embryonic yolk sac; ps: embryonic pericardial sac. Scale bar, 10 µm.

Similar EV-like structures are detected at the surface of yolk sac cells. Some appear to be internalized within yolk sac cells. Additionally, vesicle-like structures are observed on the yolk-facing side of yolk sac cells.

To investigate the distribution of maternally derived substances at the embryonic–maternal interface, immunofluorescence staining was performed following intraperitoneal injection of 2,000 kDa FITC-dextran or PBS into pregnant females.

At the follicle–yolk sac interface, punctate FITC-positive signals are detected on the embryonic side of follicular cells (Fig. 3B). These signals are smaller than 1 µm in diameter and smaller than adjacent nuclei. Similar punctate signals are detected within nonblood cells of the embryonic yolk sac. No comparable signals are observed in negative controls (IgG primary antibody or PBS-injected females).

At the follicle–pericardial sac interface, FITC-positive puncta are observed in both maternal follicular tissue and pericardial sac cells (Fig. 3C). In contrast, at the follicle–embryonic trunk boundary, FITC-positive signals are detected in the follicle but are absent from embryonic epidermal cells (*SI Appendix*, Fig. S4).

To assess whether pharmacological inhibitors affect maternal-to-embryonic transfer, pregnant females were treated with calpeptin, Y-27632, GW4869, or DMSO prior to FITC-dextran injection (*SI Appendix*, Fig. S5). Fluorescent signals are detected in embryos from all treatment groups, and quantitative analysis of fluorescence intensity in the trophic sacs showed no significant differences between embryos from DMSO-treated females and those from females treated with calpeptin, Y-27632, or GW4869.. The proportion of embryos exhibiting detectable fluorescence is 100% in the DMSO (n = 9), calpeptin (n = 10), and Y-27632 (n = 8) groups and 87.5% in the GW4869 group. Immunostaining at the follicle–yolk sac interface reveals punctate FITC signals in both follicular and yolk sac tissues in all treatment groups (*SI Appendix*, Fig. S6).

### Embryonic-to-maternal transfer of extracellular vesicles in pregnant guppies

To examine whether embryonic-to-maternal transfer of substances occurs, FITC-dextran or PBS was microinjected into the embryonic yolk sac within isolated follicles in vitro (Fig. 4A). Immunofluorescence staining was performed to assess the distribution of FITC-dextran.

**Figure 4.**
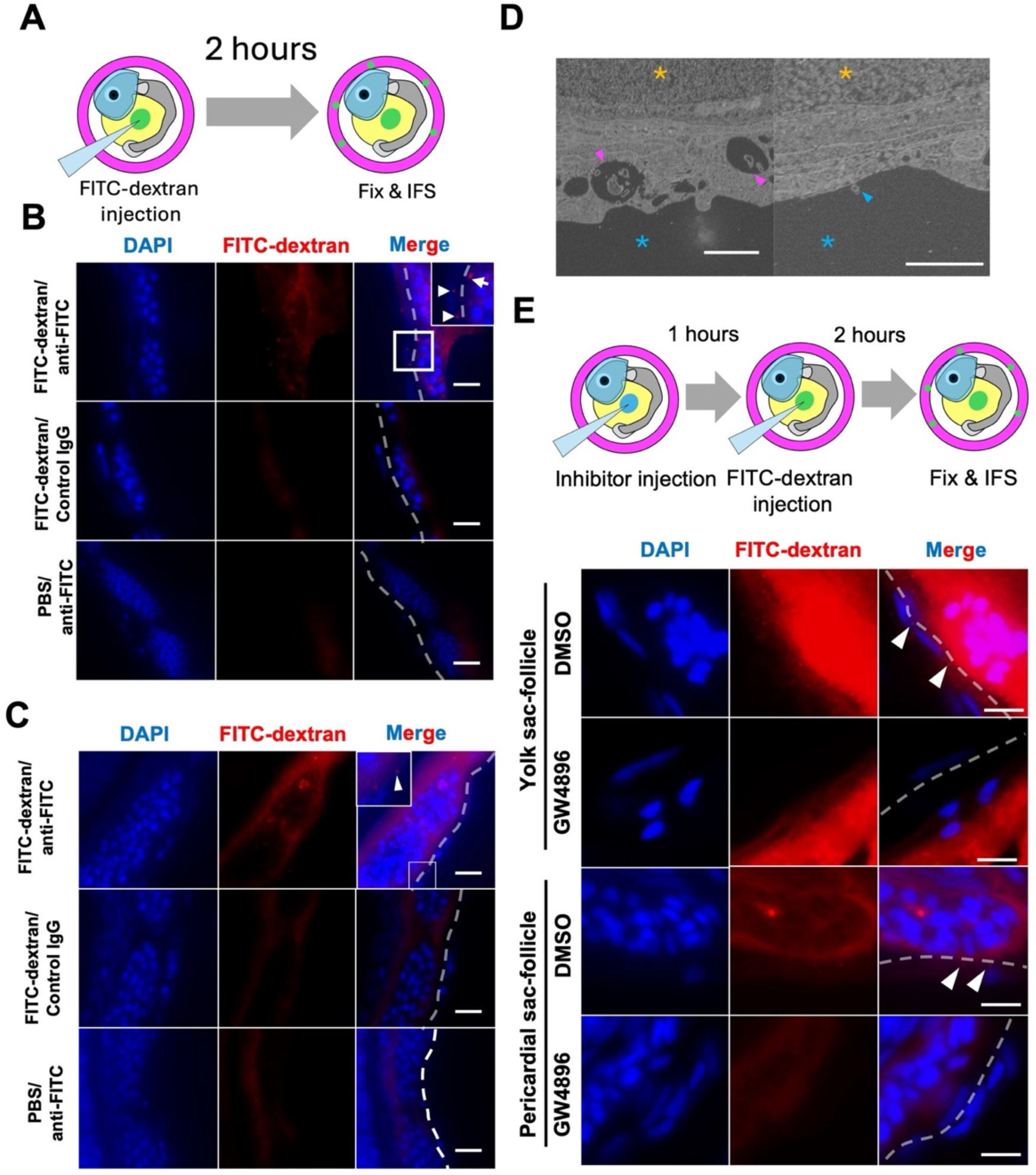
Embryonic-to-maternal material transport via exosomes.(A) Schematic illustration of the experimental system used to examine material transport from embryo to mother. FITC–dextran was injected into the embryo within the follicle *in vitro*, followed by 2 h of culture, after which the follicle containing the embryo was subjected to immunofluorescence staining. **(B, C)** Immunofluorescence staining was performed to examine whether 2,000 kDa FITC-dextran injected into the embryo within the follicle was transported from the embryonic yolk sac **(B)** or pericardial sac **(C)** to the maternal follicle. The upper row shows immunofluorescence staining using an anti-FITC antibody as the primary antibody on embryos dissected from follicles after embryonic injection of FITC-dextran. The middle row shows staining of the same specimens using rabbit IgG as a primary antibody control. The bottom row shows immunofluorescence staining using an anti-FITC antibody on embryos within follicles after embryonic injection of PBS. Enlarged panels show magnified views of merged FITC-dextran/anti-FITC images highlighting regions in which signal localization is confirmed. White arrowheads indicate FITC-dextran transferred into the maternal follicle, whereas white arrows indicate FITC-dextran within the yolk sac or pericardial sac. White dotted lines indicate the boundary between the maternal follicle and the yolk sac or pericardial sac. f: maternal follicle; ys: embryonic yolk sac; ps: embryonic pericardial sac. Scale bar, 10 µm. **(D)** Scanning electron microscopy of ultrathin sections of the yolk sac from a mid-gestation embryo. The two left panels show ultrastructural features of the maternal follicle, and the two right panels show those of the embryonic yolk sac. Blue asterisks indicate the follicular cavity in which the follicle contacts the yolk sac, and yellow asterisks indicate the lumen of blood vessels within the yolk sac. Blue arrowheads indicate exosomes secreted from the yolk sac, and magenta arrowheads indicate multivesicular bodies within yolk sac cells. Scale bar, 500 nm. **(E)** Inhibition of embryonic-to-maternal transport using an exosome inhibitor. The upper schematic illustrates the *in vitro* experimental design in which the inhibitor and FITC–dextran were administered to the embryo. The lower panels show immunofluorescence staining results demonstrating the transport of 2,000 kDa FITC-dextran from the embryonic yolk sac (top panels) or pericardial sac (bottom panels) to the maternal follicle following embryonic injection of DMSO (control, n = 8) or GW4869 (exosome inhibitor, n = 11). The first row in each set shows the follicle–embryonic interface after DMSO treatment, and the second row shows the interface after GW4869 treatment. White arrowheads indicate FITC-dextran detected within the maternal follicle, whereas white arrows indicate FITC-dextran within the yolk sac or pericardial sac. White dotted lines indicate the boundary between the maternal follicle and the yolk sac or pericardial sac. f: maternal follicle; ys: embryonic yolk sac; ps: embryonic pericardial sac. Scale bar, 10 µm.

In embryos injected with FITC-dextran, fluorescence is detected throughout the yolk sac (Fig. 4B). Punctate FITC-positive signals smaller than the nucleus are observed in outer yolk sac cells adjacent to the follicle. Similar punctate signals are detected in maternal follicular cells. No comparable signals are observed in control samples (IgG primary antibody or PBS-injected embryos).

At the pericardial sac–follicle interface, punctate FITC-positive signals are detected in pericardial sac cells and in adjacent follicular cells (Fig. 4C). These signals are absent in negative controls.

In contrast, at the follicle–embryonic trunk boundary, FITC-positive signals are absent from both of embryonic epidermal cells and maternal follicle (*SI Appendix*, Fig. S7).

To investigate ultrastructural features of the yolk sac, ultrathin sections were examined by electron microscopy (Fig. 4D). Multivesicular body–like structures are observed near the apical surface of yolk sac cells contacting the follicle. Vesicle-like structures approximately 80 nm in diameter are also observed at the cell surface.

To evaluate whether pharmacological inhibition affects embryonic-to-maternal transfer, GW4869 or DMSO was microinjected into the embryonic yolk sac prior to FITC-dextran injection (Fig. 4E). In DMSO-treated controls, punctate FITC signals are detected at the yolk sac–follicle interface in 100% of cases. In GW4869-treated samples, punctate signals are detected in 3/11 cases (27%), whereas no detectable signal is observed in 8/11 cases (73%).

These results show that embryonic injection of FITC-dextran leads to the appearance of punctate FITC-positive signals in adjacent maternal follicular tissue and that this appearance is reduced following GW4869 treatment.

## Discussion

In this study, we investigated whether maternal–embryonic substance transfer occurs between the mother and embryo in guppies (*Poecilia reticulata*), a lecithotrophic fish of the family *Poeciliidae*. Histological analysis revealed that in pregnant guppy ovaries, the maternal follicle and the embryonic pericardial sac and yolk sac constitute a maternal–embryonic interface resembling that observed in placentotrophic placentas (Fig. 1). Furthermore, black ink injection and tracking of FITC-dextran demonstrate that ovarian vascular structure and physiology change during pregnancy (Figs. 1 and 2) and that physiological mechanisms exist for both maternal-to-embryonic and embryonic-to-maternal transport (Figs. 2–4). Results from immunofluorescence staining and electron microscopy suggest that extracellular vesicles play a crucial role in maternal–embryonic substance exchange (Fig. 3) and indicate that extracellular vesicles are transported from the embryonic yolk sac and pericardial sac to maternal follicles (Fig. 4). Based on these findings, we hypothesize that during pregnancy in guppies, ovarian vascular permeability increases, allowing substances extravasated from blood vessels to be absorbed by maternal follicular cells and secreted into the follicular cavity as extracellular vesicles. These extracellular vesicles are subsequently absorbed by the embryonic yolk sac and pericardial sac, thereby facilitating nutrient transport from mother to embryo (Fig. 5). Similarly, we hypothesize that extracellular vesicles are transported from the embryonic yolk sac into the ovarian follicle, where they are taken up by maternal follicular cells, thereby enabling bidirectional maternal–embryonic transport (Fig. 5). In summary, these findings indicate that guppies possess a functional placenta in which extracellular vesicles mediate substance exchange between the maternal follicle and the embryonic yolk sac and pericardial sac.

**Figure 5.**
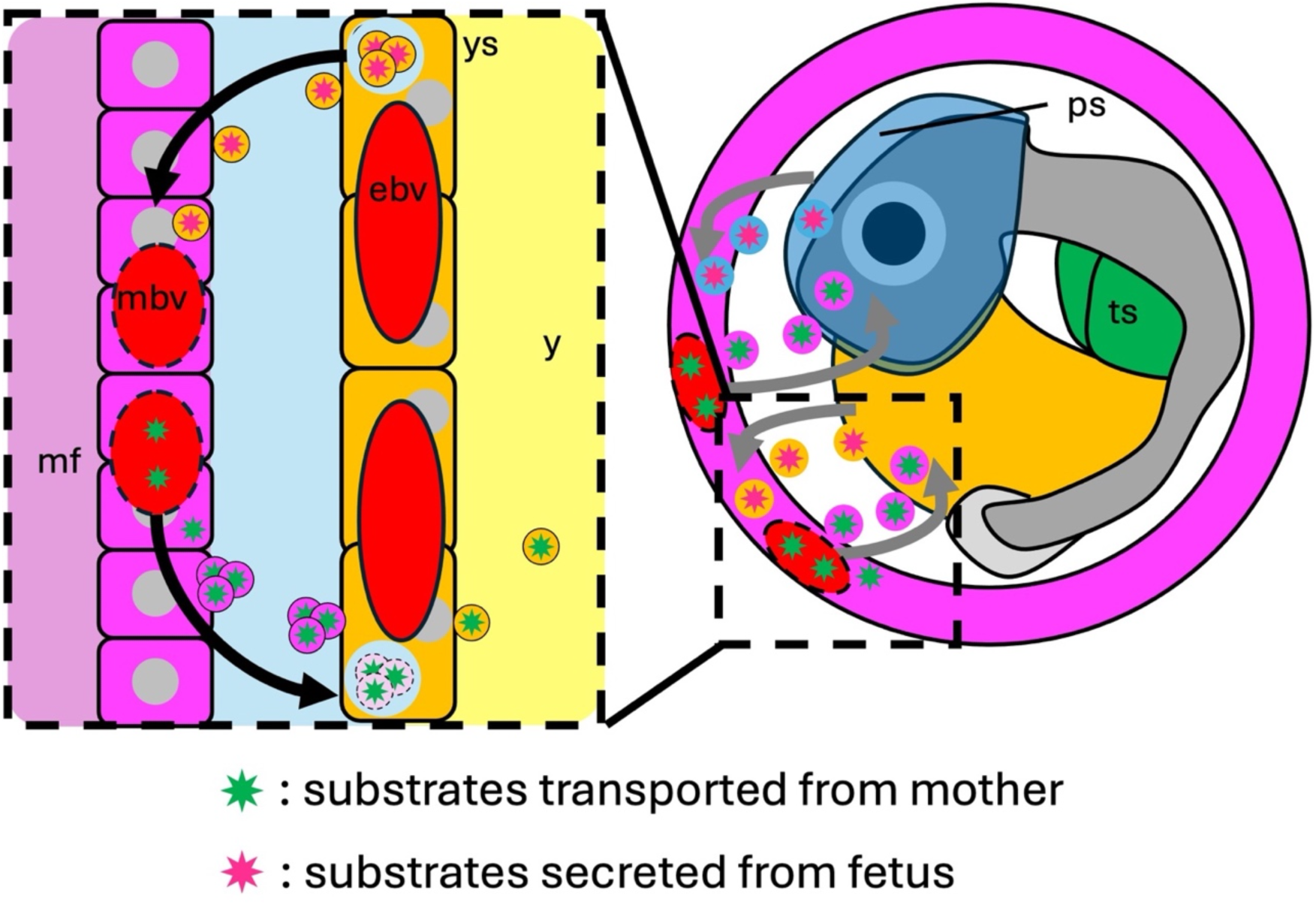
Model of maternal–embryonic transport in pregnant guppies. Schematic model illustrating bidirectional material transport between the mother and embryo in pregnant guppies. During pregnancy, vascular permeability increases in the maternal ovary, allowing substances in the maternal bloodstream to leak into the ovarian tissue. These substances are taken up by follicular cells and subsequently secreted into the follicular cavity as clusters of extracellular vesicles. The secreted extracellular vesicle clusters are internalized by endocytosis in cells of the embryonic pericardial sac and yolk sac within the follicle, and the absorbed substances ultimately accumulate in the embryonic trophic sac. Conversely, exosomes are secreted from the embryonic pericardial sac and yolk sac into the follicular cavity and are transported to the maternal follicle, indicating embryonic-to-maternal material transfer. ebv: embryonicl blood vessels; mbv: maternal blood vessels; mf: maternal follicle; ps: pericardial sac; ts: trophic sac; y: yolk; ys: yolk sac.

Visualization of the ovarian vasculature by black ink injection demonstrates that during pregnancy in guppies, the follicular vasculature undergoes pronounced remodeling from a dense, mesh-like capillary network to a hierarchically organized branched architecture (Fig. 1C). Similar remodeling of vascular networks has been reported during embryonic development and tumor angiogenesis. These changes are regulated by angiogenic factors, including vascular endothelial growth factor (VEGF), in concert with hemodynamic forces such as blood flow–derived shear stress, which together influence vascular architecture and blood perfusion(20–22). Therefore, the transition of vascular networks in maternal follicles is consistent with functional reorganization of the vascular bed, likely reflecting the increased hemodynamic demands of the pregnant ovary. In parallel, the observed leakage of high-molecular-weight (2,000 kDa) FITC-dextran into the interfollicular spaces provides evidence that vascular permeability is markedly elevated during pregnancy (Fig. 2C). Increased vascular permeability, mediated by loosening of endothelial cell–cell junctions and formation of transendothelial pores, has been reported to facilitate efficient exchange of nutrients, gases, and other small molecules within tissues(23, 24). Moreover, in humans and rodents, previous studies have shown that during early pregnancy, the maternal uterine vasculature undergoes extensive remodeling in response to angiogenic factors such as VEGF, angiopoietin, and Notch signaling, and that this remodeling is associated with increased vascular permeability(25). Together, these findings suggest that increased vascular branching enhances blood flow and perfusion within the ovary, thereby increasing the availability of oxygen and circulating nutrients, and that elevated vascular permeability allows blood-borne components to extravasate into the ovarian cavity, facilitating their subsequent transport to developing embryos within the follicles.

However, the molecular mechanisms underlying these vascular changes remain to be elucidated. Angiogenic signaling pathways, including VEGF ligands and their receptors, represent strong candidates for regulating both vascular remodeling and increased permeability, given their well-established roles in angiogenesis and endothelial function in other vertebrates. Future studies aimed at characterizing the spatial and temporal expression of these angiogenic factors, as well as experimentally testing their roles in the pregnant ovary, will be essential for understanding how vascular dynamics contribute to maternal–embryonic interactions in guppies. Therefore, identifying the angiogenic factors that regulate follicular angiogenesis and increased ovarian vascular permeability in the pregnant ovary of guppies and comparing these factors with those operating in mammals will be crucial for understanding the significance of changes in maternal vascular characteristics during embryonic development.

Furthermore, the present study demonstrates that substance transport occurs at a maternal–embryonic interface composed of the embryonic yolk sac, pericardial sac, and maternal follicle during pregnancy, thereby providing functional evidence for the presence of a placenta in lecithotrophic fish, namely guppies. Anatomical observations and histological analyses confirm that both the yolk sac and the pericardial sac persist throughout embryonic development, positioning these embryonic tissues as key components of the maternal–embryonic interface. Importantly, tissue-level observations reveal that embryonic tissues directly form the interface with the maternal follicle without an intervening space, and no direct intercellular adhesion between follicular cells and embryonic tissues is detected (Fig. 1D). These structural features are consistent with previous reports in both lecithotrophic and matrotrophic poeciliid fishes, indicating that such an interface represents a conserved architectural framework within the family.

Previous studies in Poeciliidae have proposed that the maternal follicle, yolk sac, pericardial sac, and embryo collectively contribute to maternal–embryonic substance exchange(15, 26–29). In addition, extracellular vesicles have been implicated as mediators of this transpor(30)t.

Injection of FITC-dextran into pregnant females demonstrated that maternally derived macromolecules can reach the embryo and accumulate in the trophic sac (Fig. 2). In vitro culture experiments further revealed that the route of dextran uptake by embryos differs depending on the presence of the maternal follicle (*SI Appendix*, Fig. S3). When embryos were cultured without the follicle, fluorescence was detected in the mouth, gills, and digestive tract, suggesting uptake through the oral route. In contrast, embryos cultured with the follicle intact showed dextran signals in the trophic sac, consistent with the distribution observed following maternal injection.

These observations suggest that the maternal follicle plays a key role in mediating the transfer of maternal substances to the embryo. The absence of dextran signals in the trophic sac when embryos were cultured without the follicle further indicates that trophic sac uptake is not a passive process but may require interaction with maternal follicular tissues.

Together, these findings suggest that the maternal follicle regulates the pathway by which embryos acquire external substances during pregnancy. This follicle-mediated transport mechanism may represent a specialized physiological adaptation that facilitates maternal–embryonic exchange in Poeciliidae. These findings highlight a previously unrecognized role of the maternal follicle as a functional interface for maternal–embryonic exchange in guppies.

Our immunofluorescence analyses refine this model by demonstrating that transferred substances are not delivered directly to the embryonic trunk (*SI Appendix*, Fig. S4). Instead, they are selectively taken up by the yolk sac and pericardial sac via extracellular vesicles originating from the maternal follicle (Fig. 3B,C). These findings suggest that the yolk sac and pericardial sac function as primary reception and processing sites for maternally derived substances, rather than the embryo serving as the initial target.

Taken together, these results support the idea that a common vesicle-mediated transport mechanism operates at the maternal–embryonic interface in poeciliid fishes, irrespective of whether the species is predominantly lecithotrophic or matrotrophic. Such a mechanism may represent an evolutionarily conserved strategy for maternal provisioning that predates or operates independently of more specialized placental structures.

Nevertheless, alternative or complementary transport pathways should also be considered. In matrotrophic poeciliids, expression of certain transporter genes in the maternal follicle and embryonic yolk sac has been documented(15), raising the possibility that small molecules may be transferred directly via membrane transporters in addition to extracellular vesicle–mediated delivery. Future studies integrating molecular, ultrastructural, and functional approaches will be required to clarify the relative contributions of these pathways and to determine how different modes of maternal provisioning are coordinated during pregnancy.

FITC–dextran injected into pregnant females localizes predominantly within a membranous tissue (trophic sac) situated in the embryonic body cavity (Fig. 2E). To date, detailed knowledge of internal organogenesis in poeciliid embryos remains limited. Although gross anatomical descriptions of embryonic internal organs at successive developmental stages have been reported for mosquitofish(19), corresponding histological analyses are lacking. As a result, the organ homologous to the trophic sac has previously been misidentified as the swim bladder.

By integrating anatomical observations of guppy embryos with histological analyses of embryonic tissue sections, the present study demonstrates that the trophic sac is a distinct organ separate from the swim bladder, which is characterized by the presence of gas gland cells (Fig. 2F). This distinction clarifies a long-standing ambiguity in poeciliid embryonic anatomy. Furthermore, the observation that maternally injected FITC-dextran accumulates within the trophic sac suggests that this organ functions as a storage site for substances transferred from the mother during pregnancy.

In addition, immunofluorescence staining reveals that a portion of maternally derived FITC-dextran localizes to and is secreted on the yolk-facing side of the yolk sac (Fig. 3B, yellow panel). Consistent with this observation, electron microscopy demonstrates that yolk sac cells secrete extracellular vesicles toward the yolk sac lumen and that part of the trophic sac is in direct contact with the yolk sac (Fig. 3A). Together, these findings suggest a stepwise transport pathway in which substances transferred from the mother to the yolk sac are subsequently packaged into extracellular vesicles and transported from the yolk sac to the trophic sac.

Despite these insights, the molecular composition of the substances accumulated within the trophic sac remains unknown. Determining whether the trophic sac primarily stores nutrients, signaling molecules, or other maternally derived factors will be essential for understanding the functional significance of this organ in embryonic development.

Notably, no internal organ comparable to the trophic sac has been described in developing embryos of other teleost fishes, suggesting that the trophic sac represents a lineage-specific innovation. Although the present study provides functional and anatomical evidence for the trophic sac, it does not address its developmental origin, including its germ layer derivation or morphological changes during embryogenesis. Future studies examining the developmental mechanisms underlying trophic sac formation are therefore expected to shed light on the evolutionary emergence of maternal nutrient provisioning strategies in the Poeciliidae lineage.

Fluorescent immunostaining and electron microscopy analyses indicate that maternal-to-embryonic substance transport in guppies is mediated by extracellular vesicles. Consistent with this observation, vesicle-like structures have previously been reported in the ovarian follicles of the viviparous poeciliid *Heterandaria formosa*(30), suggesting that vesicle-mediated transport may represent a conserved feature within the family Poeciliidae. Despite this, the physiological mechanisms underlying the formation and release of these follicle-derived vesicles remain unresolved.

In the present study, pharmacological inhibition of pathways associated with microvesicle or exosome biogenesis does not suppress maternal–embryonic transport (*SI Appendix*, Figs. S5 and S6), implying that the extracellular vesicles observed here are unlikely to be generated through canonical microvesicle- or exosome-dependent pathways. Furthermore, ultrastructural analyses reveal that vesicles secreted from the maternal follicle are characterized by a highly electron-dense membrane and are released in clustered aggregates (Fig. 3A). These features differ from those typically described for exosomes or microvesicles(31–33). Notably, morphologically similar clustered exosome-like vesicles have been reported during lens development(34), raising the possibility that alternative vesicle biogenesis pathways operate in specific developmental or physiological contexts.

Taken together, these findings suggest that the extracellular vesicles secreted from maternal follicles in guppies may represent a distinct class of vesicles, separate from classical exosomes or microvesicles. However, definitive classification requires further investigation. Elucidating the cellular origin, molecular composition, and biogenetic pathways of these vesicles will be essential for understanding their physiological roles and clarifying how vesicle-mediated transport contributes to maternal provisioning during pregnancy in poeciliid fishes.

In contrast, immunofluorescence staining demonstrates that substance transport from the embryo to the mother is also mediated by extracellular vesicles (Fig. 4B,C). Consistent with this observation, electron microscopy reveals the presence of multivesicular bodies within yolk sac cells (Fig. 4D), supporting the involvement of an exosome-derived vesicle secretion pathway. Notably, administration of the exosome biogenesis inhibitor GW4869 to the embryo significantly inhibits embryonic-to-maternal transport (Fig. 4E), whereas maternal-to-embryonic transport remains unaffected (*SI Appendix*, Figs. S5 and S6). These findings strongly suggest that substance transfer from the embryo to the mother occurs through exosome secretion, whereas transport in the opposite direction is regulated by a distinct physiological mechanism.

Such directionality in vesicle-mediated transport is reminiscent of mammalian placentation, in which exosomes secreted by fetal trophoblast cells are released into the maternal circulation and function in immune modulation, metabolic regulation, and intercellular signaling(35–37). Although the present study does not directly investigate the functions or molecular cargo of exosomes secreted from guppy embryos, the phenomenon of fetal-to-maternal exosome transport appears to be conserved across distantly related viviparous vertebrates. These observations suggest that exosome-mediated signaling from embryo to mother may represent a fundamental and evolutionarily conserved feature of placental interactions.

The differential effects of GW4869 on the two transport directions further indicate that maternal-to-embryonic and embryonic-to-maternal exchanges are governed by independent molecular mechanisms. Such separation may be crucial for preventing interference between opposing transport systems operating at the maternal–embryonic interface. While exosome-associated molecules are likely to play a central role in embryonic-to-maternal transport, the physiological mechanisms underlying extracellular vesicle formation and transfer from the maternal follicle to the embryo remain unresolved. Nevertheless, ultrastructural observations suggest that uptake into yolk sac cells occurs via endocytosis (Fig. 3A), implicating the involvement of endocytosis-related genes and pathways in maternal-to-embryonic transport.

In addition, the present study does not determine the specific substances contained within the extracellular vesicles secreted by either the maternal follicle or the embryo. It is plausible that these vesicles transport nutrients from the mother, thereby fulfilling a placental function. Indeed, in the family Poeciliidae, large proteins such as vitellogenin have been shown to be secreted from the maternal ovary and absorbed by a pseudoplacental structure known as the trophotaeniae(9). Beyond nutrient provision, extracellular vesicles are also known to carry signaling molecules, including proteins and microRNAs (miRNAs), which can induce specific physiological responses in recipient tissues(38, 39). Such signaling functions may therefore be essential for coordinating normal pregnancy in guppies.

Taken together, these findings suggest that during pregnancy in guppies, extracellular vesicles may transport not only nutritional components such as vitellogenin but also regulatory molecules such as miRNAs, thereby facilitating both metabolic exchange and intercellular communication between mother and embryo. Furthermore, previous studies in Poeciliidae have reported that expression levels of vesicle-associated genes in maternal follicles fluctuate in response to the matrotrophic index(15). This observation implies that vesicle-mediated transport mechanisms—including cargo composition and transport capacity—may vary substantially among species within the family. Consequently, evolutionary changes in genes associated with extracellular vesicle biogenesis, secretion, and uptake may underlie interspecific variation in maternal provisioning strategies and contribute to the evolution of matrotrophy within the Poeciliidae.

This study demonstrates that in guppies, maternal–embryonic exchange of substances occurs via extracellular vesicles at a maternal–embryonic interface that lacks direct tissue adhesion. Such a nonadhesive maternal–embryonic topology appears to be common among poeciliid fishes(15, 26–29), suggesting that vesicle-mediated maternal–embryonic exchange is a shared and conserved feature within this family.

In contrast, although placental morphology and maternal–fetal interactions vary widely among vertebrates, cellular adhesion between maternal and fetal tissues is generally observed in most vertebrate placentas. In eutherian mammals such as mice and humans, the fetal mesoderm-derived chorionic sac forms a placenta in association with the maternal endometrium, with fetal tissues invading the maternal decidua(2, 3). In other placental mammals, including horses and pigs, fetal tissues adhere to maternal tissues without invasion(2, 3). Outside mammals, marsupials and *Mabuya* lizards form placentas involving both the fetal chorionic sac and the endoderm-derived yolk sac in contact with the endometrium(6), whereas in hammerhead sharks, the fetal yolk sac adheres directly to the uterine epithelium to establish a placental connection(40). On the other hand, previous studies have shown that in the porcine placenta, maternally derived uteroferrin is absorbed not only at sites of adhesion between maternal and fetal tissues but also within fetal areolae that are not directly attached to maternal tissue(41). These findings suggest that substance transport can occur at maternal–fetal interfaces even in the absence of direct cellular adhesion. The present study shows that in guppies, the maternal follicle does not adhere to embryonic tissues, including the yolk sac and pericardial sac (Fig. 1D). Instead, extracellular vesicles are secreted from the maternal follicle and transported to these embryonic tissues (Fig. 2), enabling maternal–embryonic substance exchange in the absence of tissue adhesion. These findings reveal a fundamentally different mode of maternal–fetal interaction in which vesicle-mediated transport substitutes for direct cellular contact.

Taken together, these results indicate that guppies employ a maternal provisioning strategy that is distinct from the placental systems described in mammals and other viviparous vertebrates. Vesicle-mediated transport across a nonadhesive maternal–fetal interface may represent an alternative evolutionary solution for sustaining pregnancy, thereby highlighting the remarkable diversity of placental organization and maternal–fetal exchange mechanisms among vertebrates.

## Materials and Methods

### Ethical statement

This study was approved by the Animal Care Committee of Meiji University (approval number: MUIACUC2024-09). In accordance with the institutional guidelines, a minimal number of animals were euthanized by chilling on ice at 4 °C.

### Fish

Guppies (*Poecilia reticulata*) were purchased from Ichigaya Fish Center (Tokyo, Japan) and Reimix (Nagoya, Japan). Guppies were maintained at 27 °C under a 14/10 h light/dark photoperiod. Adults were fed brine shrimp and Hikari Peret Guppy (Kyorin Co., Ltd., Himeji, Japan), whereas fry were fed brine shrimp.

### Anatomy

Pregnant female guppies were anesthetized in ice-cold water, and their ovaries were removed. The ovaries were placed in phosphate-buffered saline (PBS), and embryos were extracted at the developmental stage equivalent to Ph29 of *Gambusia affinis*(*19*) by rupturing the follicles with fine tweezers. The extracted embryos were photographed using a stereomicroscope (SMZ800N; Nikon, Tokyo, Japan).

Next, both nonpregnant and pregnant females were anesthetized in ice-cold water, and their caudal fins were photographed using the SMZ800N. Subsequently, 100 µL of black ink (Sumi no Hoshi; Kuretake Co., Ltd., Nara, Japan) was injected into the abdominal cavity. The females were washed three times with PBS and maintained in 0.3% seawater for 1 h. They were then reanesthetized, and their caudal fins were photographed using the SMZ800N. After observation of the caudal fins, the ovaries were removed, washed three times with PBS, and fixed overnight in 4% paraformaldehyde (PFA) in PBS. The ovaries were subsequently washed three times for 5 min each in PBS, and the follicular vasculature was photographed using the SMZ800N.

### Histology

Pregnant females were anesthetized, and their ovaries were removed. The ovaries were washed with PBS and then fixed overnight in Davidson’s fixative at 4 °C. The ovaries were washed three times for 1 h each in 70% ethanol and then maintained overnight at 4 °C in 70% ethanol. Meanwhile, embryos were removed from the ovaries, washed with PBS, and fixed overnight in 4% PFA in PBS at 4 °C. The embryos were washed three times for 5 min each in PBS and then maintained overnight at 4 °C in 70% ethanol. The specimens were dehydrated and embedded in paraffin using an automated fixation and embedding device (RH-12DM; Sakura Finetek, Tokyo, Japan). Paraffin blocks were sectioned at 5 µm, mounted on glass slides (Muto Chemical, Tokyo, Japan), and incubated overnight at 40 °C. The specimens were deparaffinized, rehydrated, and stained with hematoxylin and eosin (HE). The specimens were sealed with Entellan New (Merck, Sigma-Aldrich, St. Louis, MO, USA) and observed using a BZ-X710 microscope (Keyence, Osaka, Japan).

### FITC-dextran injection

Pregnant females were anesthetized in ice-cold water and injected intraperitoneally with either 100 µL of 1 µg/mL 2,000 kDa fluorescein isothiocyanate (FITC)-dextran (Sigma-Aldrich) in PBS or 100 µL of PBS alone. After injection, the fish were washed three times with PBS and maintained in 0.3% seawater. At 1 h and 24 h after injection, the fish were anesthetized, and their caudal fins were observed using a fluorescence stereomicroscope (MVX10; Olympus, Tokyo, Japan). After fluorescence observation of the caudal fins at 24 h post-injection, the ovaries were removed, washed three times with PBS, and observed using the MVX10. Embryos were then removed from each ovary using forceps, washed three times with PBS, and observed using a fluorescence microscope. For selected embryos observed under the fluorescence microscope, the yolk sac was ruptured with forceps after initial observation, the yolk was removed, the embryos were washed with PBS, and they were observed again using the MVX10.

### In vitro assay of maternal-to-embryonic transport

Pregnant females were anesthetized, and their ovaries were removed. After washing the ovaries in PBS, they were divided into two halves along the left–right axis using forceps in Leibovitz’s L-15 medium with L-glutamine with L-glutamine (Gibco, Thermo Fisher Scientific, Waltham, MA, USA), taking care not to rupture the follicles. Half of the collected embryos had their maternal follicles removed using forceps. Embryos with follicles removed and embryos with follicles intact were cultured separately for 2 h at room temperature in Leibovitz’s L-15 medium containing 1 mg/ml 2000 kDa FITC-dextran. After incubation, follicles were removed from intact samples. The yolk was removed by rupturing the yolk sac with forceps, and embryos were washed three times with PBS before fluorescence observation using an MVX10.

### Immunofluorescent staining

Pregnant females were anesthetized and injected intraperitoneally with either 100 µL of 1 µg/mL 2,000 kDa FITC-dextran in PBS or 100 µL of PBS alone. After injection, the fish were washed three times with PBS and maintained in 0.3% seawater. Twenty-four hours after injection, the ovaries were removed, washed three times with PBS, and fixed overnight in 4% PFA in PBS. The fixed ovaries were washed three times for 5 min each in PBS and then sequentially immersed in 15% and 30% sucrose in PBS for 1 h each, followed by overnight immersion in a solution of 30% optimal cutting temperature (OCT) compound in 30% sucrose in PBS. The ovaries were embedded in OCT compound, sectioned at 10 µm using a cryomicrotome, mounted on silane-coated slides, and air-dried overnight at 37 °C. The slides were washed three times for 5 min each in PBS and then blocked with 1% bovine serum albumin in PBS at room temperature for 30 min. Subsequently, the sections were incubated with primary antibodies at 4 °C for 24 h using either a 1:100 dilution of rabbit polyclonal anti-FITC antibody (Invitrogen, Thermo Fisher Scientific, Carlsbad, CA, USA; #71-1900) or a 1:500 dilution of rabbit polyclonal anti-mouse immunoglobulin G (IgG) antibody (Sigma-Aldrich, A9044). After washing three times for 5 min each in PBS, the sections were incubated with a secondary antibody at 4 °C for 2 h using a 1:500 dilution of goat polyclonal anti-rabbit IgG antibody (Abcam, Cambridge, UK; ab150080) in 0.1% Triton X-100 in PBS. After washing three times for 5 min each in PBS, the slides were mounted with 4′,6-diamidino-2-phenylindole (DAPI) diluted 1:1,000 in 50% glycerol in PBS and observed using a BZ-X710 microscope (Keyence, Osaka, Japan).

### Inhibition of maternal-to-embryonic transport

Inhibitor concentrations were determined based on previous studies(42–45). Calpeptin (catalog no. 7396; Selleckchem, Houston, TX, USA), Y-27632 (MedChemExpress, NJ, USA; HY-10583), and GW4869 (MedChemExpress; HY-19363) were dissolved in dimethyl sulfoxide (DMSO) to prepare stock solutions at concentrations of 72 mg/mL, 64 mg/mL, and 0.2 mg/mL, respectively. Prior to injection, the stock solutions were diluted with PBS as follows: calpeptin was diluted 14-fold, Y-27632 was diluted 600-fold, and GW4869 was diluted 10-fold.

Adult guppies were anesthetized by immersion in ice-cold water, and 100 µL of each inhibitor solution (calpeptin in DMSO and PBS, Y-27632 in DMSO and PBS, or GW4869 in DMSO and PBS) or control solution (10% DMSO in PBS) was injected into the body cavity using a 27-gauge needle. Two hours after inhibitor administration, 100 µL of 2,000 kDa FITC-dextran dissolved in PBS, or PBS alone, was injected intracoelomically.

Following a 5 h incubation period, guppies were reanesthetized, and embryos were dissected for fluorescence observation using an MVX10 stereomicroscope (Olympus, Tokyo, Japan). In addition, ovaries were collected from females treated under identical conditions and processed for immunofluorescence staining using an anti-FITC antibody.

### Electron microscopic observation

Pregnant females were anesthetized in ice-cold water, and the ovaries were removed. The ovaries were washed with PBS and fixed overnight in 2.5% glutaraldehyde and 2% PFA in PBS. After washing three times with PBS, the follicles and embryos were separated from the ovaries using forceps. The yolk sacs were then separated from the embryos using forceps.

The separated follicles and yolk sacs were sequentially dehydrated in 50%, 70%, 80%, 90%, 95%, and 100% ethanol at room temperature for 10 min each. The specimens were then immersed in acetone, followed by sequential infiltration with 50% epoxy resin (Nisshin EM Quetol-812) in acetone and 100% epoxy resin at room temperature for 1 h each. The yolk sacs and follicles were embedded in epoxy resin and polymerized at 60 °C for 60 h. Polymerized samples were sectioned at 100 nm using an ultramicrotome and stained with uranyl acetate substitute (UraniLess; Micro to Nano) and lead citrate (Micro to Nano). Backscattered electron images were obtained using an SU9000 field-emission scanning electron microscope (Hitachi, Tokyo, Japan) at 1 kV.

### Embryo-to-mother transport

Pregnant females were anesthetized, and their ovaries were removed. After washing the ovaries in PBS, they were divided into two halves along the left–right axis using forceps in Leibovitz’s L-15 medium with L-glutamine, taking care not to rupture the follicles. Each divided follicle was placed in separate Leibovitz’s L-15 medium. Using a glass capillary, 0.5 µL of either 1 µg/mL FITC-dextran in PBS or PBS alone was injected into the yolk sac of the embryo within each follicle. The follicles were then washed three times with PBS, placed in Leibovitz’s L-15 medium, and cultured at room temperature for 2 h. After culture, the follicles were washed three times with PBS, fixed overnight in 4% PFA in PBS, and subjected to immunofluorescence staining, as described for the ovaries of females injected with FITC-dextran.

### Inhibition of transport from embryo to mother

Pregnant females were anesthetized, and the ovaries were dissected and immediately washed in PBS. Individual follicles were carefully isolated and longitudinally bisected using fine forceps in Leibovitz’s L-15 medium, taking care not to rupture the follicular structure. Each half follicle was transferred to separate dishes containing Leibovitz’s L-15 medium.

Using a glass capillary, 0.5 µL of either 0.2 mg/mL GW4869 dissolved in DMSO or DMSO alone was injected into the yolk sac of the embryo within each follicle. One hour after inhibitor injection, 0.5 µL of 1 µg/mL FITC–dextran in PBS, or PBS alone, was injected into the yolk sac using a glass capillary.

Following injection, the follicles were washed three times with PBS and cultured in Leibovitz’s L-15 medium at room temperature for 2 h. After culture, the follicles were washed three additional times with PBS, fixed overnight in 4% PFA in PBS, and processed for immunofluorescence staining as described for ovaries obtained from females injected with FITC-dextran.

## Acknowledgments

We thank the Research Promotion Projects for Young Researchers at Meiji University for supporting this work. We are grateful to Makoto Sugiyama for insightful discussions. We also thank the High-Voltage Electron Microscope Laboratory (HVEM), Institute of Materials and Systems for Sustainability (IMaSS), Nagoya University, Tokai National Higher Education and Research System (THERS), for support in providing devices for ultrastructural analysis.

## Figures

**Figure S1.**
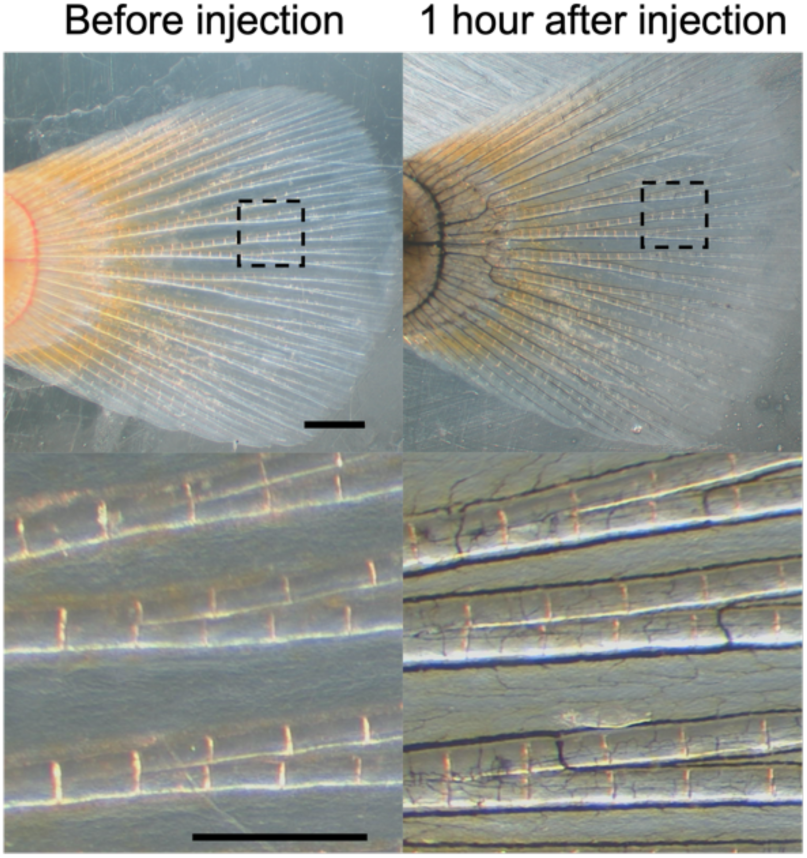
Visualization of maternal blood vessels following black ink injection. The caudal fin of a pregnant female was observed under a stereomicroscope before (left) and 1 h after (right) intraperitoneal injection of black ink into the same individual. The upper panels show low-magnification views of the entire anal fin, whereas the lower panels show high-magnification views of the regions outlined by dotted lines in the upper panels. Scale bars, 1 mm (upper panels) and 500 µm (lower panels).

**Figure S2.**
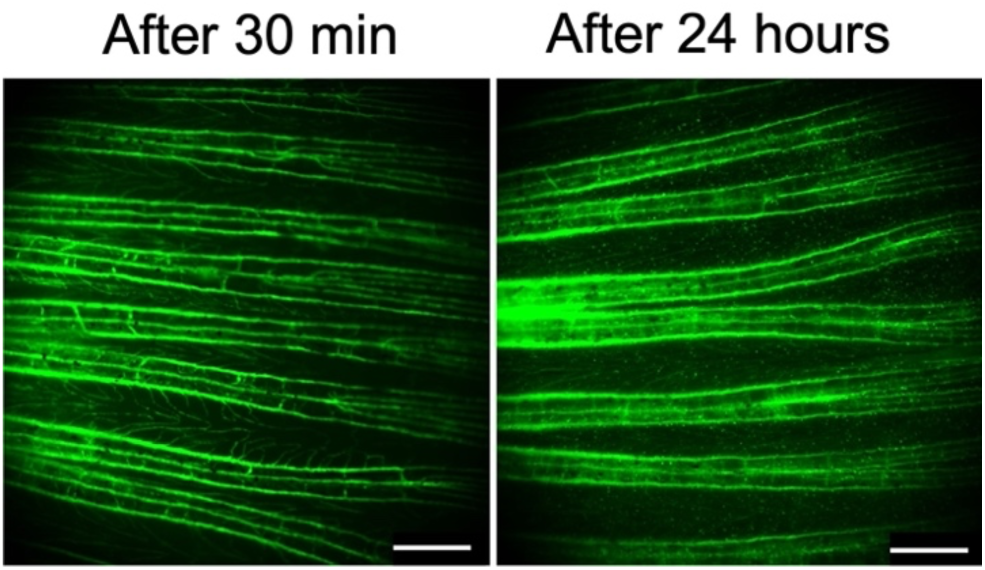
Confirmation of tail fin vascular morphology and permeability by FITC–dextran injection. Vascular permeability in the tail fins of pregnant females was examined following intraperitoneal injection of fluorescent dextran. The left panel shows tail fin vasculature observed under a fluorescence stereomicroscope 30 min after injection of 2,000 kDa FITC-dextran. The right panel shows the same individual 24 h after injection. Only minimal leakage of FITC-dextran was detected in the fin membrane. Scale bar, 500 µm.

**Figure S3.**
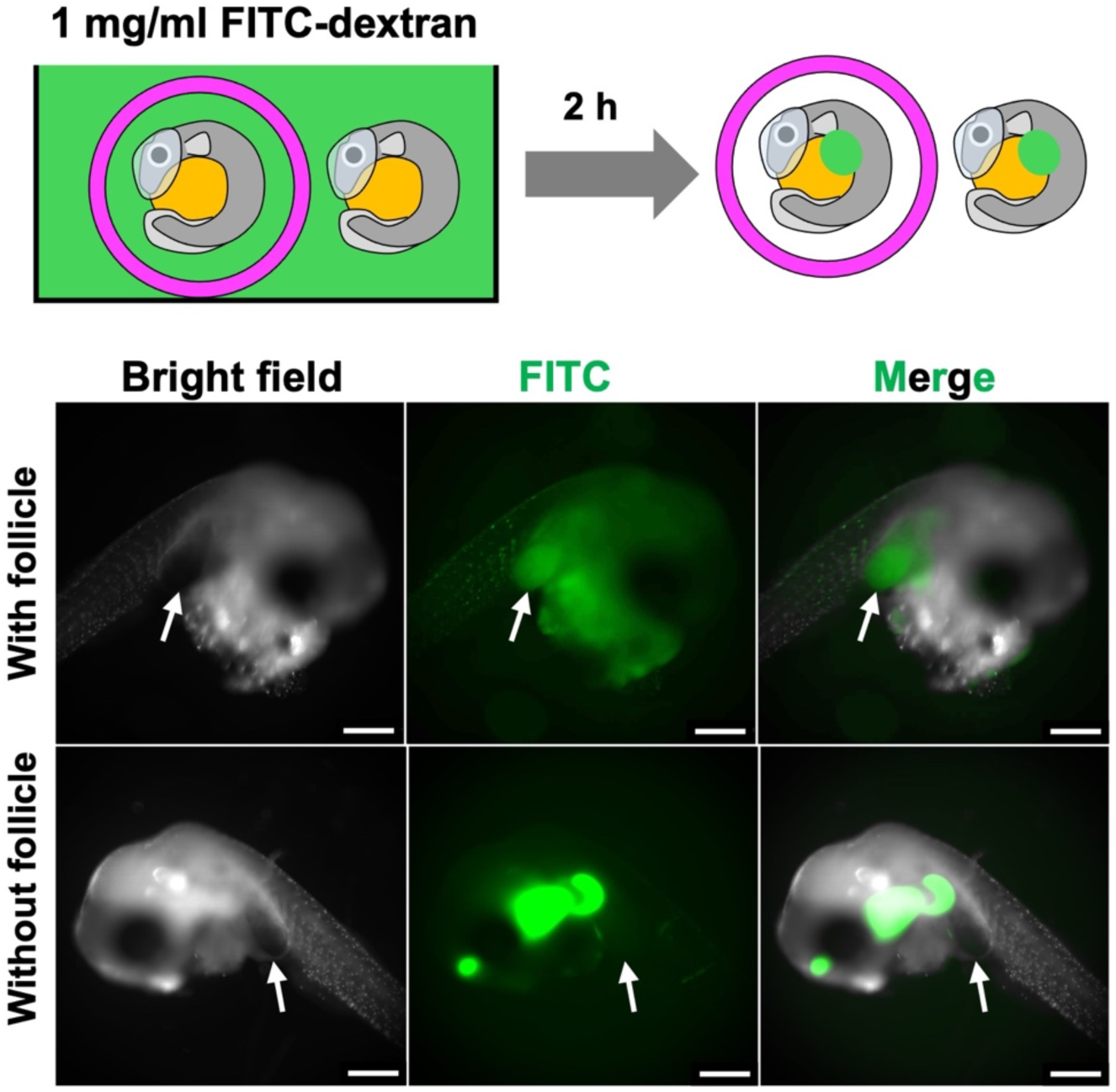
FITC-dextran uptake by embryos with or without the maternal follicle. Embryos with the maternal follicle intact or after follicle removal were cultured in medium containing 2000 kDa FITC-dextran to examine dextran uptake by embryonic tissues. The schematic (upper panel) shows the experimental design. Images (lower panel) show embryos after 2 h of culture following yolk removal. Upper images show embryos cultured with the maternal follicle intact, whereas lower images show embryos after follicle removal. White arrows indicate trophic sacs. Scale bar, 500 µm.

**Figure S4.**
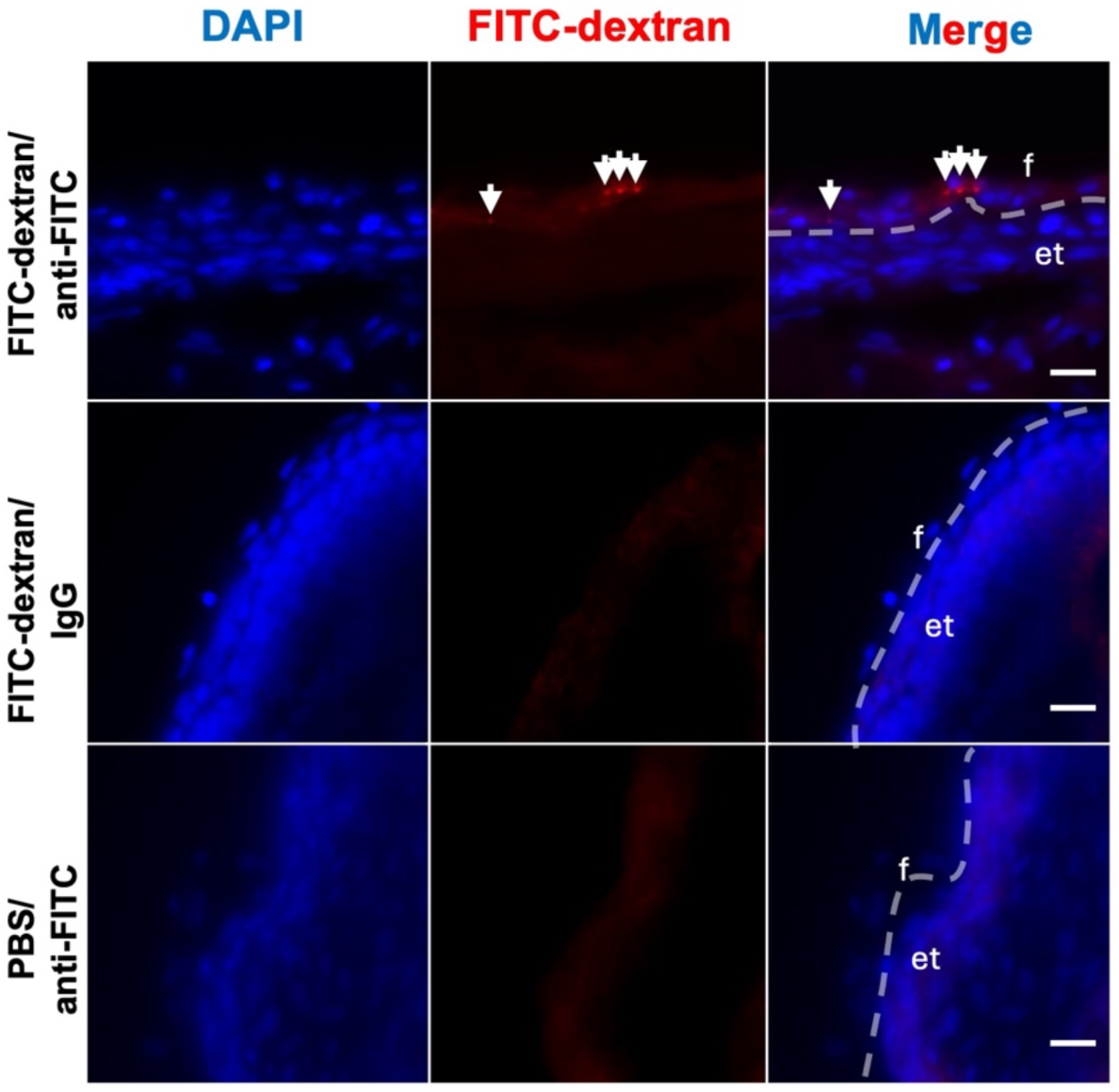
Interface between the embryonic trunk and the maternal follicle. To examine whether 2,000 kDa FITC-dextran injected into pregnant females is directly transported from the maternal follicle to the embryonic trunk, immunofluorescence staining was performed using an anti-FITC antibody. The upper panel shows immunofluorescence staining of a embryo within a follicle dissected from a female injected with FITC-dextran. The middle panel shows staining of the same specimen using rabbit IgG as a primary antibody control. The bottom panel shows immunofluorescence staining using an anti-FITC antibody on embryos within follicles dissected from females injected with PBS. Blue signals indicate DAPI-labeled nuclei, and red signals indicate FITC-dextran detected by immunofluorescence. White dotted lines indicate the boundary between the maternal follicle and the embryonic trunk, and white arrows indicate detected FITC-dextran signals. f: maternal follicle; et: embryonic trunk. Scale bar, 10 µm.

**Figure S5.**
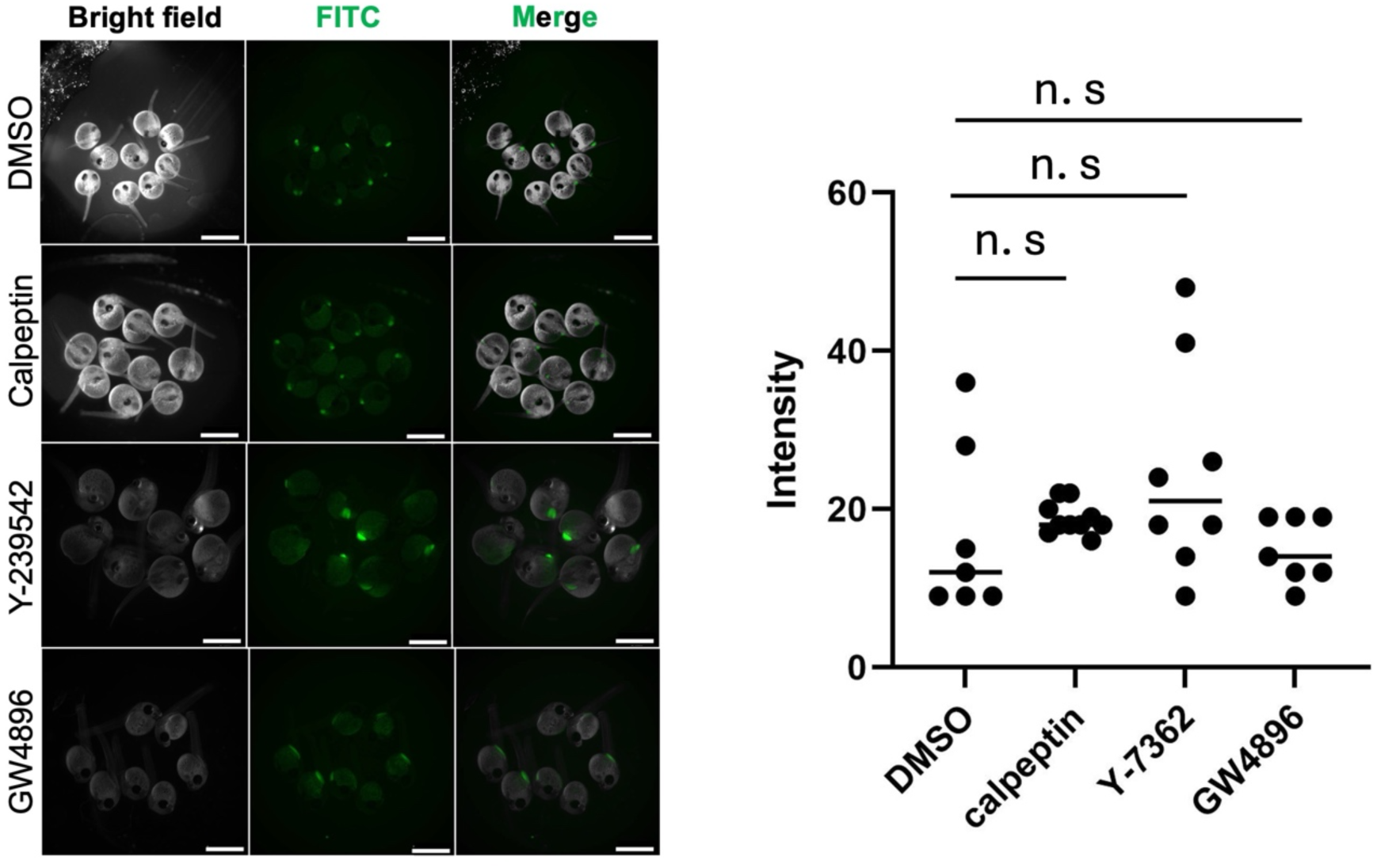
Inhibitors targeting microvesicle and exosome formation do not inhibit maternal-to-embryonic transport of FITC–dextran. The left panels show representative fluorescence stereomicroscope images of embryos dissected from pregnant females injected with inhibitors followed by 2,000 kDa FITC-dextran. The first row shows embryos from females injected with DMSO (control) followed by FITC-dextran. The second and third rows show embryos from females injected with the microvesicle formation inhibitors calpeptin (second row) or Y-27632 (third row), followed by FITC-dextran. The fourth row shows embryos from females injected with GW4869, an inhibitor of exosome formation, followed by FITC-dextran. Scale bar, 2 mm. The right panel shows quantification of the maximum fluorescence intensity detected in the embryos shown in the left panels. An unpaired two-tailed Student’s *t*-test was conducted, and no significant differences were observed among treatment groups (n.s.).

**Figure S6.**
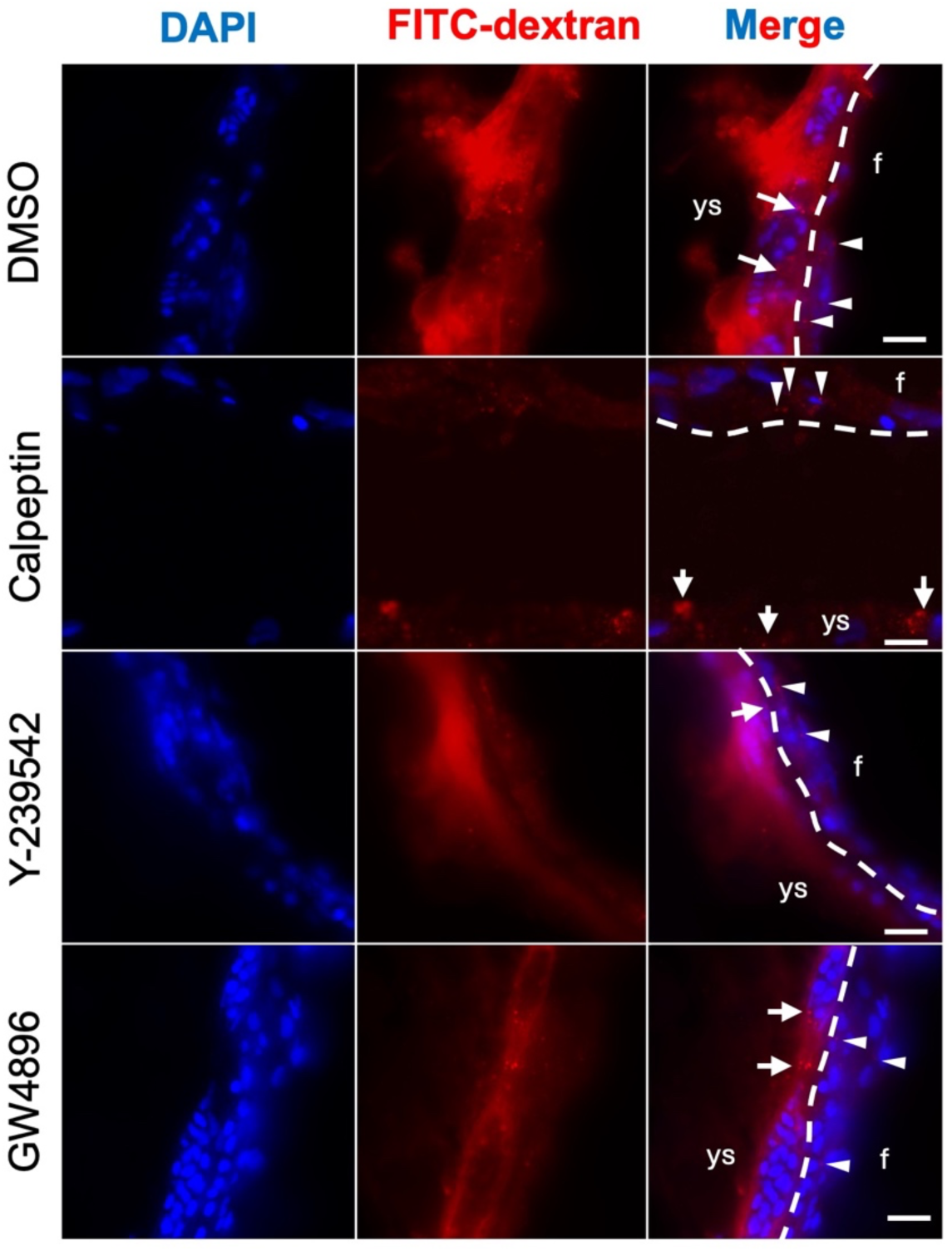
Inhibitors of microvesicle and exosome formation do not block extracellular vesicle–mediated transport from mother to embryo. To examine whether inhibition of microvesicle or exosome formation affects maternal-to-embryonic transport, pregnant females were injected with inhibitors followed by 2,000 kDa FITC–dextran, and immunofluorescence staining using an anti-FITC antibody was performed on embryos within isolated follicles. The first panel shows embryos from follicles dissected from females injected with DMSO (control) followed by FITC-dextran. The second and third panels show embryos from follicles dissected from females injected with calpeptin (second panel) or Y-27632 (third panel), inhibitors of microvesicle formation, followed by FITC-dextran injection. The fourth panel shows embryos from follicles dissected from females injected with GW4869, an inhibitor of exosome formation, followed by FITC-dextran injection. Blue signals indicate DAPI-labeled nuclei, and red signals indicate FITC-dextran detected by immunofluorescence. White dotted lines indicate the boundary between the maternal follicle and the embryonic trunk. White arrows indicate FITC-dextran signals detected in the maternal follicle, and white arrowheads indicate FITC-dextran signals detected in the yolk sac. f: maternal follicle; ys: embryonic yolk sac. Scale bar, 10 µm.

**Figure S7.**
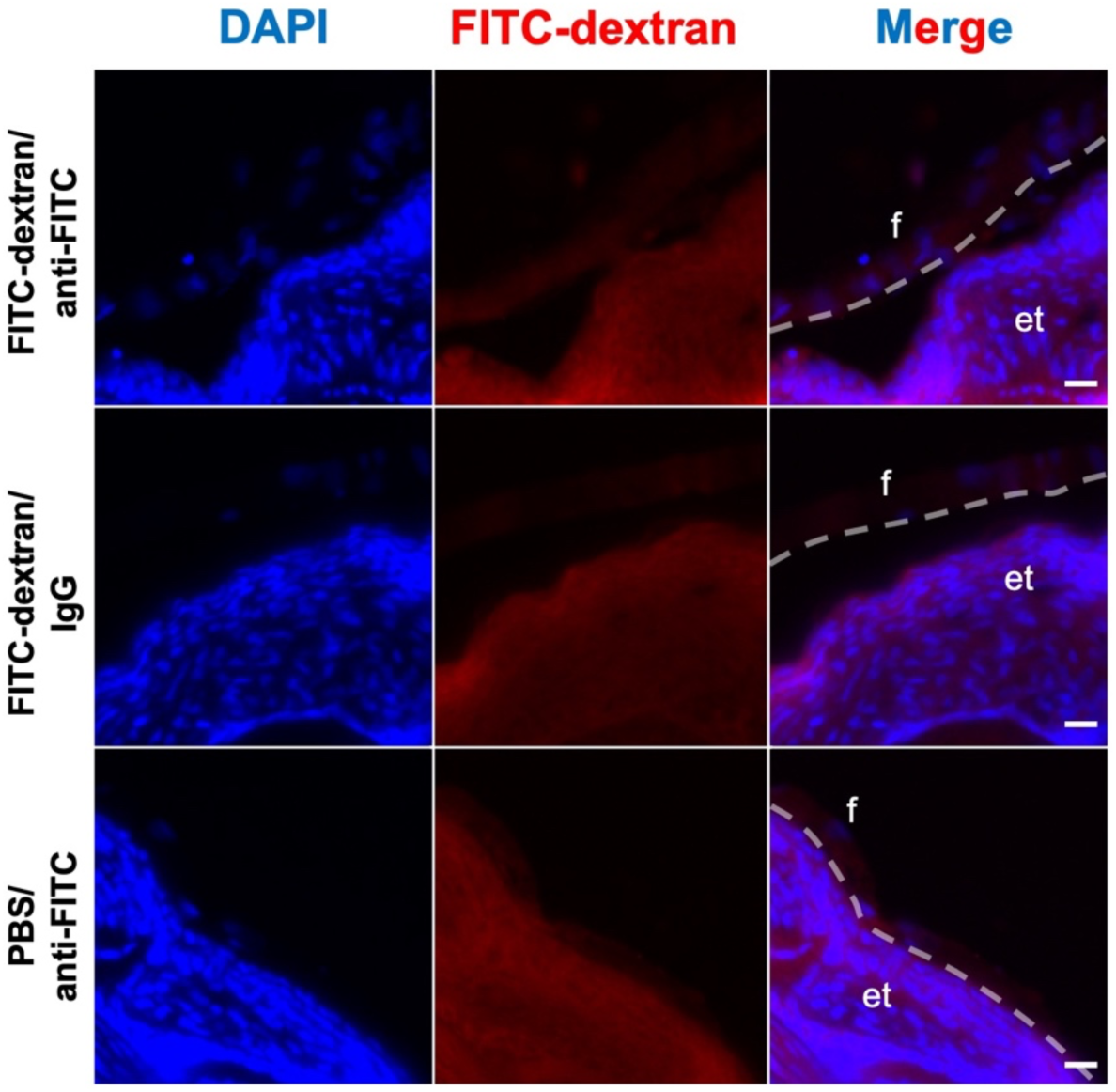
Absence of detectable transport from the embryonic trunk to the maternal follicle at the embryonic trunk–follicle interface. To examine whether 2,000 kDa FITC–dextran injected into the embryo within the follicle is transported from the embryonic trunk to the maternal follicle, immunofluorescence staining using an anti-FITC antibody was performed. The top panel shows immunofluorescence staining of a embryo injected with FITC-dextran within the follicle. The middle panel shows staining of the same specimen using rabbit IgG as a primary antibody control. The bottom panel shows immunofluorescence staining using an anti-FITC antibody on a embryo injected with PBS within the follicle. Blue signals indicate DAPI-labeled nuclei, and red signals indicate FITC-dextran detected by immunofluorescence. White dotted lines indicate the boundary between the maternal follicle and the embryonic trunk. White arrows indicate detected FITC-dextran signals. No FITC-dextran signal was observed within the maternal follicle under these conditions. f: maternal follicle; et: embryonic trunk. Scale bar, 10 µm.

**Movie S1.**
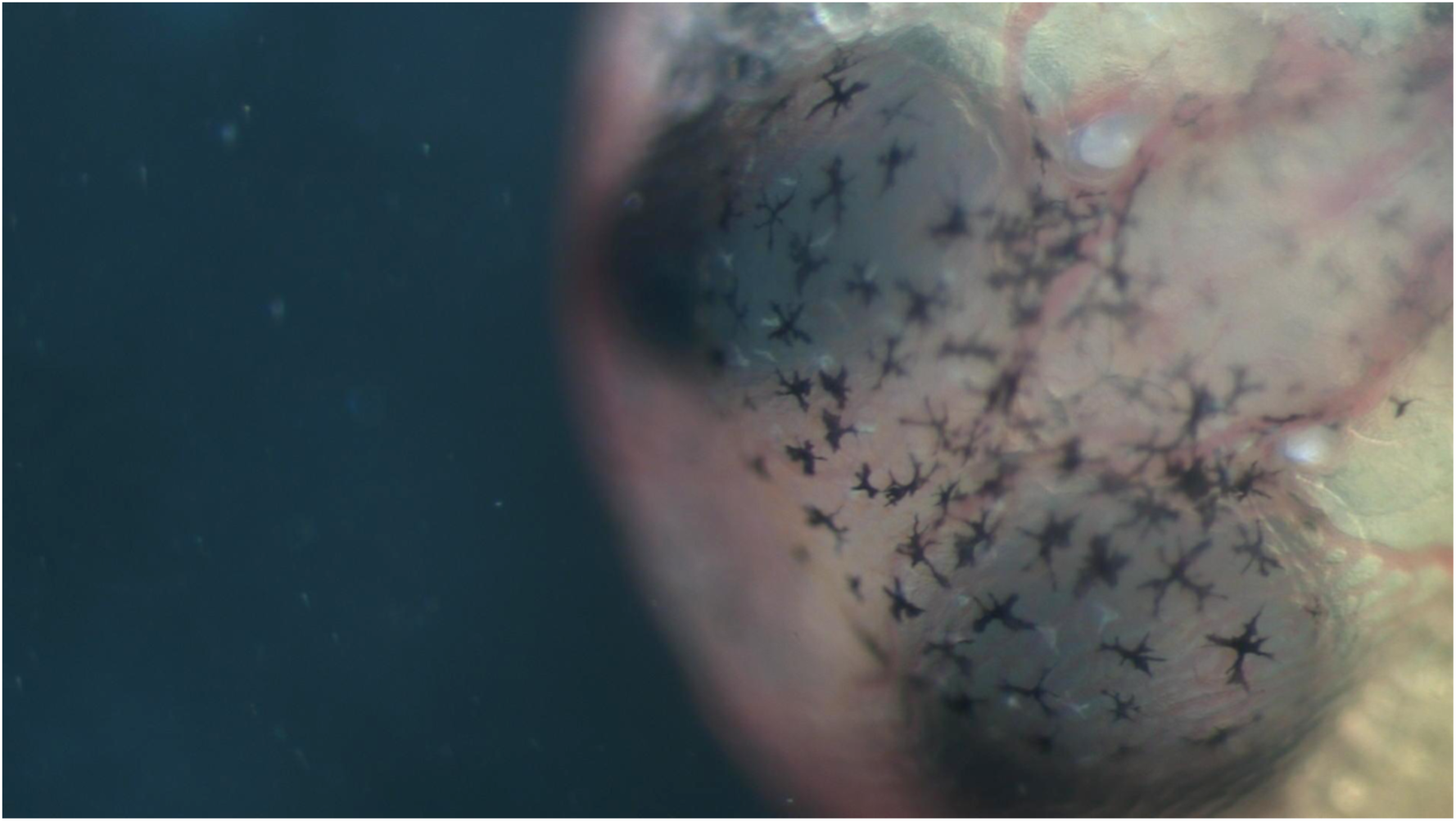
Blood flow in the pericardial sac. A viable embryo was gently removed from the maternal follicle and transferred into a dish containing PBS. Blood flow within the pericardial sac was recorded at room temperature (27 °C) using a stereomicroscope. The dimensions of the video field are 1,855 µm × 1,043 µm.

**Movie S2.**
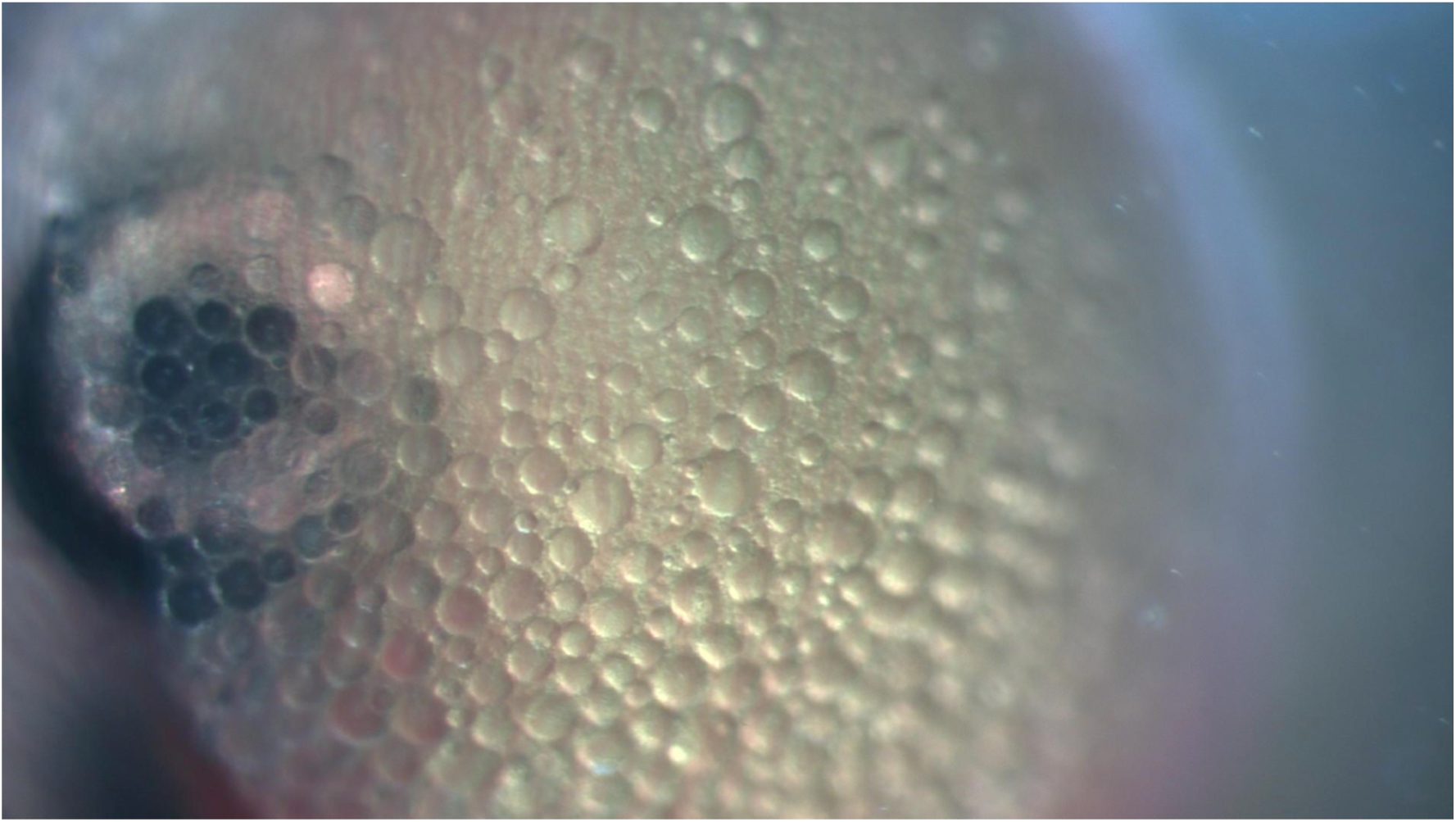
Blood flow in the yolk sac. A viable embryo was gently removed from the maternal follicle and transferred into a dish containing PBS. Blood flow within the yolk sac was recorded at room temperature (27 °C) using a stereomicroscope. The dimensions of the video field are 1,855 µm × 1,043 µm.

**Movie S3.**
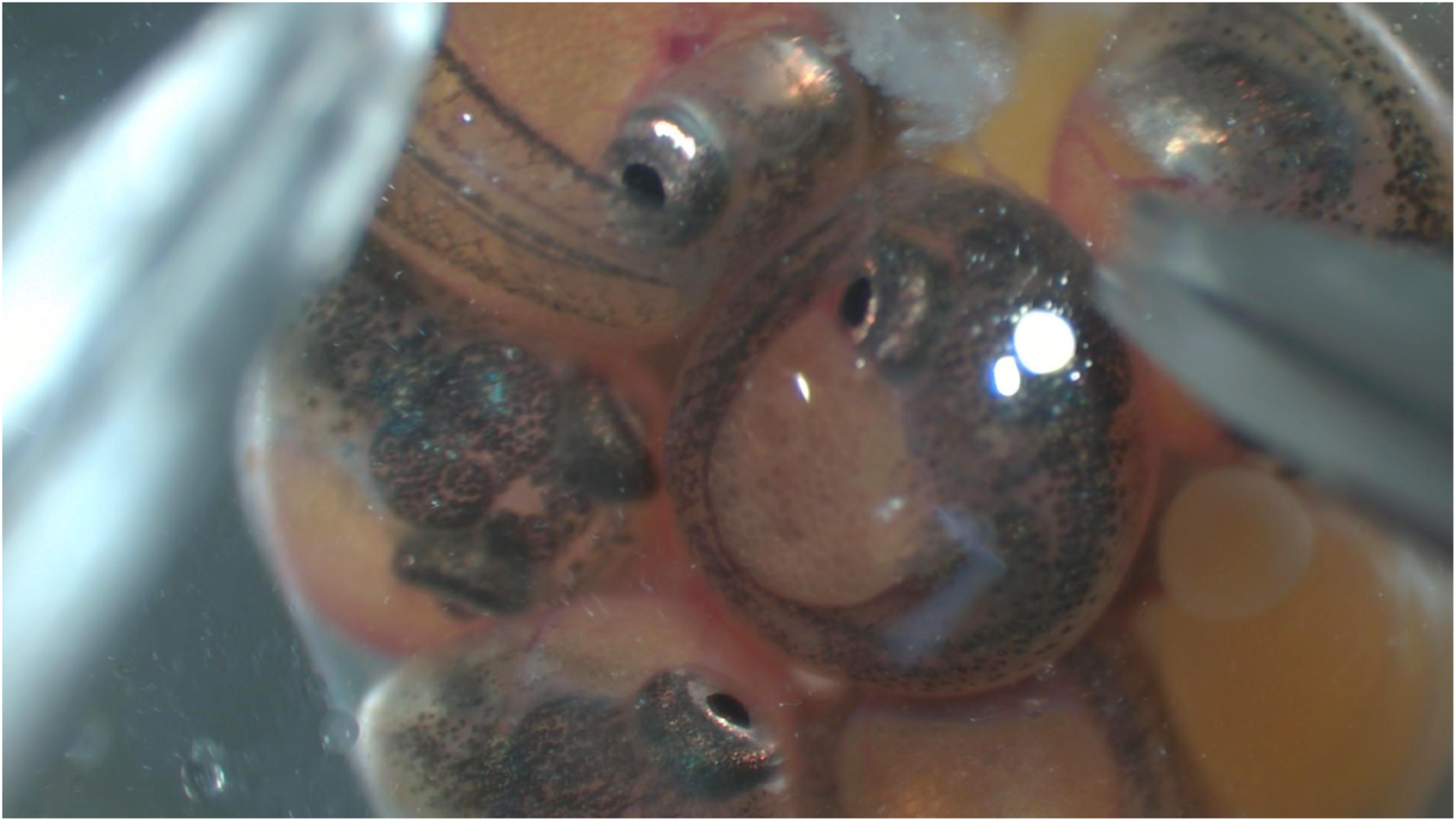
Embryonic movement within the ovary during late pregnancy. Ovaries were dissected from late-pregnant females and placed in a dish containing PBS. The ovaries were recorded at room temperature (27 °C) using a stereomicroscope. To induce embryonic movement, embryos within the follicles were gently stimulated with fine forceps. The dimensions of the video field are 4,637 µm × 2,609 µm.

